# QUANTITATIVE PROTEOMIC REVEALS HEME OXYGENASE-1 LOCALIZATION TO CELL-SURFACE LIPID RAFTS AND ITS ASSOCIATION WITH FERROPORTIN IN IRON-LOADED MACROPHAGES

**DOI:** 10.64898/2026.09.14.750975

**Authors:** Alexandra Willemetz, Anne Auriac, Christian Federici, Lorenne Robert, Marjorie Leduc, Luc Camoin, François Canonne-Hergaux

## Abstract

**Background:** Macrophages play a central role in systemic iron homeostasis by recycling iron from senescent erythrocytes through the only known cellular iron exporter, ferroportin (Fpn). Previous studies have shown that Fpn localizes to membrane microdomains, or lipid rafts, which are important for its function and regulation by hepcidin. Iron strongly promotes the enrichment of Fpn in lipid rafts. However, the proteomic composition of these domains during iron overload remains largely unexplored.

**Objective:** This study aimed to characterize iron-induced changes in the macrophage lipid raft proteome in order to identify potential functional partners of Fpn and better define the cellular mechanisms involved in the response to iron overload.

**Methods:** Quantitative iTRAQ-based proteomic analysis was performed on detergent-resistant membranes (DRMs) isolated by iodixanol density-gradient ultracentrifugation from J774a.1 macrophages treated with iron-NTA (FeNTA). Key proteomic observations were further validated in both J774a.1 cells and bone marrow-derived macrophages (BMDMs) using western blot, cell-surface biotinylation, and confocal immunofluorescence microscopy. The regulatory role of the transcription factor Nrf2 was also assessed using Nrf2-knockout mouse models.

**Results:** In addition of Fpn, our lipid raft proteome identified 79 proteins significatively upregulated upon iron treatment, including, antioxidant proteins like the peroxiredoxin-1 (Prdx1), the glucose-6-phosphate dehydrogenase X-linked (G6pdx), as well as heme oxygenase-1 (Hmox1). Bioinformatic and functional analyses indicated that this response is associated with nuclear translocation of Nrf2, which is required for the transcriptional induction of Fpn and Hmox1. Whereas Fpn was detected in lipid rafts under basal conditions, Hmox1 was recruited to these domains only after iron or heme treatment. Confocal microscopy and cell-surface biotinylation assays confirmed that Hmox1 and Fpn colocalize at the plasma membrane, particularly within caveolae, specialized invaginated lipid raft domains.

**Conclusions:** Iron or heme overload induces a major remodeling of the macrophage membrane, leading to the formation of specialized lipid raft platforms enriched in antioxidant and iron handling proteins. The coordinated localization of Hmox1 and Fpn at the cell surface suggests a coupled mechanism linking heme catabolism to iron export. This organization may help limit intracellular iron accumulation and protect the plasma membrane from iron-mediated lipid peroxidation.

**Graphical abstract:** 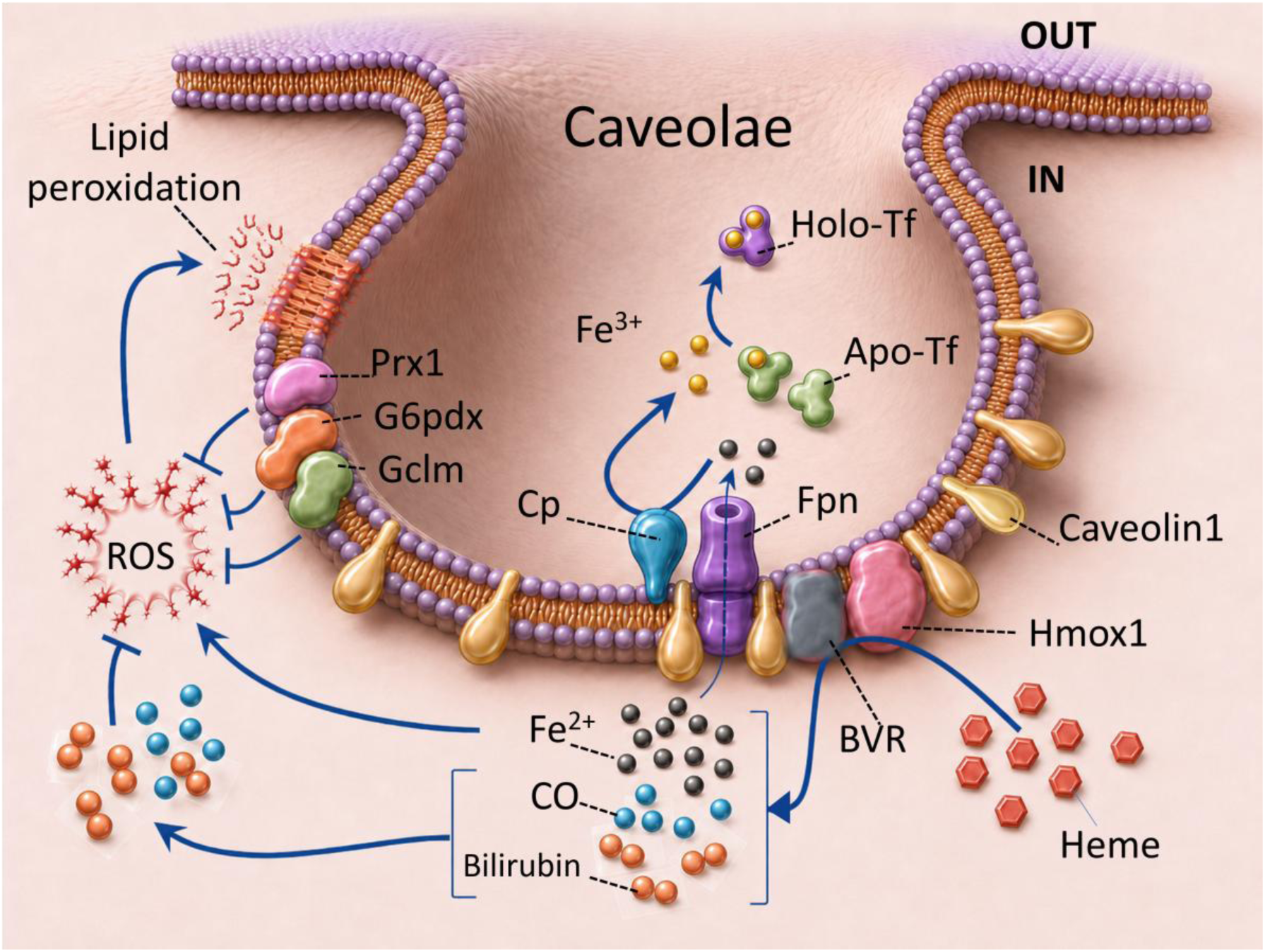

Cellular iron overload induces oxidative stress and may promote membrane lipid peroxidation. Proteomic analysis in macrophages of detergent-resistant membranes (lipid raft) enriched in the iron exporter ferroportin (Fpn) suggests that heme or iron (FeNTA) overloading promotes the recruitment of several proteins to caveolae, specialized lipid raft membrane invaginations enriched in caveolin. Components of the antioxidant response, including peroxiredoxin 1 (Prdx1), glucose-6-phosphate dehydrogenase X-linked (G6pdx), and glutamate-cysteine ligase modifier subunit (Gclm), were detected in these fractions, consistent with the establishment of a localized protective response that involves Nrf2-dependent signaling. Heme oxygenase-1 (Hmox1), the enzyme responsible for heme degradation, was also associated with the plasma membrane and colocalized with Fpn and caveolin1 at the cell surface. Hmox1 converts heme into ferrous iron, carbon monoxide (CO), and biliverdin. Biliverdin can subsequently be reduced to bilirubin by biliverdin reductase (BVR), which has been reported present in lipid rafts (Kim et al., 2004). Both CO and bilirubin have antioxidant and cytoprotective properties and may therefore limit ROS production and oxidative membrane damage. In parallel, the ferroxidase ceruloplasmin (Cp), previously shown to be present with cell surface lipid rafts (Marques et al., 2012), may facilitate iron export by oxidizing Fe²⁺ to Fe³⁺ for loading onto transferrin. This coordinated mechanism could restrict intracellular iron accumulation, limit Fenton chemistry, and protect membrane integrity during heme or iron overload. The graphical abstract was developed from a hand-drawn sketch using Microsoft PowerPoint, with image-generation support provided by ChatGPT.

## Introduction

Iron homeostasis is a fundamental biological process, and macrophages play a central role in its systemic regulation, notably by recycling iron from senescent erythrocytes (Soares & Hamza, 2016). Ferroportin (Fpn, Slc40a1) is the only known cellular iron exporter (Abboud & Haile, 2000; Donovan et al., 2000; McKie et al., 2000) and its activity at the macrophage cell surface represents therefore a critical control point for iron availability in the body (Delaby, Pilard, Gonçalves, et al., 2005; Knutson et al., 2003; Sabelli et al., 2017). Indeed, Iron recycling in macrophages thought the phagocytosis of senescent erythrocytes is a major process in systemic iron homeostasis and involved Fpn (Soares & Hamza, 2016).

Previous studies have demonstrated that Fpn is upregulated by iron in both J774a1 macrophages and bone marrow derived macrophages (BMDM), and is downregulated by hepcidin, the systemic iron regulatory hormone (Delaby, Pilard, Gonçalves, et al., 2005). Hepcidin induces rapid internalization and degradation of Fpn, at least in part through lysosomal pathways (Nemeth et al., 2004; Delaby, Pilard, Gonçalves, et al., 2005; Knutson et al., 2005). Importantly, Fpn has also been shown to be strongly enriched in membrane microdomains or lipid rafts in iron-treated macrophages (Canonne-Hergaux et al., 2006; Marques et al., 2012; Auriac et al., 2010). Lipid rafts correspond to sphingolipid and cholesterol enriched membrane domains present in cellular membrane and containing specific proteins. Organized as platform, lipid rafts play numerous roles in cell organization, signaling and homeostasis (Lingwood & Simons, 2010).

Interestingly, disruption of lipid raft integrity using cholesterol lowering drugs in macrophages impairs hepcidin-mediated regulation of Fpn, suggesting that this membrane environment and associated protein partners are required for efficient hepcidin action (Auriac et al., 2010). Depletion of cholesterol was also shown to reduce the iron export activity of Fpn (Debbiche et al., 2023). In addition, the glycosylphosphatidylinositol-anchored ceruloplasmin (GPI-Cp), the ferroxidase involved in the cellular iron efflux by Fpn, was found to be present with the DRM containing Fpn (Marques et al., 2012). More recently, we have reported, in hepcidin-deficient mouse models, the presence of Fpn in DRM from liver (hepatocytes, Kupffer cells, and liver macrophages), and spleen (splenic macrophages) but not from intestine (enterocytes) (Besson et al., 2024). Interestingly, Fpn exhibits tissue-specific isoforms across duodenal enterocytes, hepatocytes, and macrophages, driven by different localization such as lipid rafts and post-transcriptional modifications such as glycosylation patterns (Canonne-Hergaux et al., 2006). These differences could influence the iron transport activity of Fpn and its regulation, with hepcidin-induced degradation occurring more efficiently in macrophages than in enterocytes (Chaston et al., 2008). Notably, in mice, during the suckling period, enterocyte Fpn displays reduced responsiveness to circulating hepcidin and altered electrophoretic mobility, suggesting alternative splicing or distinct glycosylation compared to the Fpn forms in adults (Frazer et al., 2017).

Altogether, such observations suggest that the biochemical and the cellular environment of Fpn in biological membrane is determinant for the function and the regulation of Fpn. Despite major insights in the study of Fpn (transport activity, structural studies, hepcidin regulation…), the membrane and proteomic environment surrounding Fpn in different cells and tissues remain underexplored, requiring further investigation to fully elucidate its functional implications in iron homeostasis and related disorders.

We therefore designed a quantitative proteomic analysis of the lipid rafts or DRM compartment containing Fpn. Our goal was to document the proteomics changes in cell surface lipid rafts in iron overload in macrophages. One of our hypothesis states that the partners involved in the regulation of Fpn expression and its transport activity should also be induced by iron within the same compartment.

## Materials and Methods

### Animals or ethic statement

SWISS mice were purchased from Janvier (CS4105 Le Genest St Isle F-53,941 St Berthevin Cedex, France). C57BL6J mice (wild type) and the NFE2-related factor 2 (Nfe2rf2, alias NRF2) null mice/C57BL6J, originally developed by Masayuki Yamamoto, (University of Tsukuba; 1996), were provided by the RIKEN BRC through the National Bio-Resource Project of the MEXT, Japan. All the procedures done with animals were conducted in accordance with the Guide for the Care and Use of Laboratory Animal.

### L-Cell-Conditioned Medium (LCCM)

L929 cells (ATCC, NCTC clone 929) were cultured in RPMI-GlutaMAX supplemented with 10% heat-inactivated FCS, 2 mM L-glutamine, and antibiotics. Cells were seeded at 2.5 × 10⁴ cells/ml in 75- or 150 cm² flasks and maintained for 8 days until confluence. Supernatants were collected, filtered through 0.22 µm membranes, aliquoted, and stored at −20 °C. LCCM was used as a source of CSF1.

### Macrophage cell cultures

The mouse macrophage cell line J774a1 was obtained from ATCC (TIB-67) and cultured in DMEM, 10% FBS in Petri dishes (10 cm diameter) for protein isolation and extraction. The bone marrow derived macrophages (BMDM) were isolated from femurs of 6–8-week-old Swiss mice as previously described (Delaby al., 2005). Cells were flushed with HBSS and were cultured in RPMI-GlutaMAX supplemented with 10% heat-inactivated FCS, 10% LCCM, 2 mM L-glutamine, and antibiotics. At day 4, adherent cells were washed with HBSS–HEPES, and medium was renewed daily until day 7. Cells were seeded at 2.5 × 10⁵ cells/ml in Petri dishes for protein analysis, or at 1.25 × 10⁵ cells/ml on 10 mm glass coverslips in 24 well plates for immunofluorescence or in 96 well culture plates for In-Cell western blot (LI-COR).

### Treatments

An FeNTA (nitrilotriacetate) stock solution (10mM FeCl3 / 40 mM NTA) was prepared just prior to the experiment and dilute in the culture medium to obtain a final solution of 100 μM FeCl₃ / 400 μM NTA for BMDM or 200 μM FeCl₃ / 800 μM NTA for J774a1. Cells were then incubated overnight (O/N). For Heme treatment, NS (Normosang®; human haemin; Orphan Europe) was obtained from the Centre Français des Porphyries, Hôpital Louis Mourier (Colombes, France) and used at the concentration of 10 µM O/N.

### Cytosolic and membrane protein fractionation

BMDM or J774A.1 cells grown in Petri dishes were washed twice with 5 ml of ice-cold PBS. Cells were harvested by scraping in 2 ml PBS containing 2 mM EDTA at 4°C and collected by centrifugation for 5 min at 230 × g. Cell pellets were resuspended in lysis buffer supplemented with protease inhibitors (1/100; Sigma P8340) and PMSF (100 µg/ml) and lysed by mechanical shearing through a 26-gauge needle (20 passes). Lysates were centrifuged at 1,100 × g for 10 min at 4°C to remove nuclei, unlysed cells, and large debris. The resulting supernatant was transferred to ultracentrifuge tubes and centrifuged in a TLA55 rotor at 186,000 × g for 45 min at 4°C. The cytosolic fraction was recovered from the supernatant, whereas the membrane fraction was obtained from the pellet, which was resuspended in TNE buffer (0,01 M Tris-HCl; 0,5M EDTA, 0,1 M NaCl) supplemented with protease inhibitors (1/100; Sigma P8340) and PMSF (100 µg/ml).

### Cell surface protein isolation by biotinylation

Cell surface proteins were isolated using the Pierce Cell Surface Protein Isolation Kit (Thermo Scientific) according to the manufacturer’s instructions. J774A.1 cells were cultured in T75 flasks (four per condition), washed twice with ice-cold PBS, and incubated with biotinylation reagent for 30 min at 4°C. The reaction was quenched, and cells were harvested by scraping. Cells were then washed with TBS, lysed, and sonicated (low power). Lysates were centrifuged at 10,000 × g for 2 min at 4°C, and supernatants were incubated with NeutrAvidin agarose for 1 h at room temperature. After washing, bound proteins were eluted in sample buffer and stored at −20°C prior to western blot analysis.

### Isolation of rafts/detergent resistant membranes

J774 or BMDM culture Petri dishes were washed twice with 5 ml of cold PBS, and cells were harvested by scraping them into 2 ml of PBS containing 2 mM EDTA. A total of 10^8^ J774a1 cells and 10^7^ BMDM cells were centrifuged for 5 minutes at 230 g at 4°C. The cell pellets were resuspended in 600 μl of MBS buffer containing 1% Triton X-100, protease inhibitors (1/100), and PMSF (1/100). Cells were lysed on ice for 1 hour. To enhance lysis efficiency, mechanical shearing was performed by passing the cell lysate 20 times through a 26-gauge needle. A density gradient was prepared by adding 2.4 ml of 50% (w/v) iodixanol working solution to the cell lysate. The 50% (w/v) iodixanol working solution is obtained by mixing 5 vol of an OptiPrep™ Density Gradient Medium (60% [w/v] iodixanol; Sigma-Aldrich, D1556) with 1 vol of a buffer containing 0.25 M saccharose, 6 mM EDTA and 120 mM Tricine-NaOH pH 7,6 to the cell lysate. The cell lysate was then carefully layered at the bottom of a gradient tube (Beckman). Sequentially, 4.5 ml of 30% (w/v) iodixanol solution and 4.5 ml of 20% (w/v) iodixanol solution were gently layered on top. The 30% (w/v) and 20% (w/v) iodixanol solutions were prepared by diluting the 50% (w/v) iodixanol working solution with a buffer containing 0.25 M saccharose, 1 mM EDTA and 10 mM Tricine-NaOH pH 7,6. The gradient was then centrifuged at 260,000 g for 16 hours at 4°C in an SW41 rotor. Finally, 1 ml fractions were collected and analyzed by western blot or mass spectrometry (MS/MS).

### Proteomic analysis

Quantitative iTRAQ proteomic analysis was performed as outlined in Figure 1. Principal steps are detailed below.

**Figure 1.**
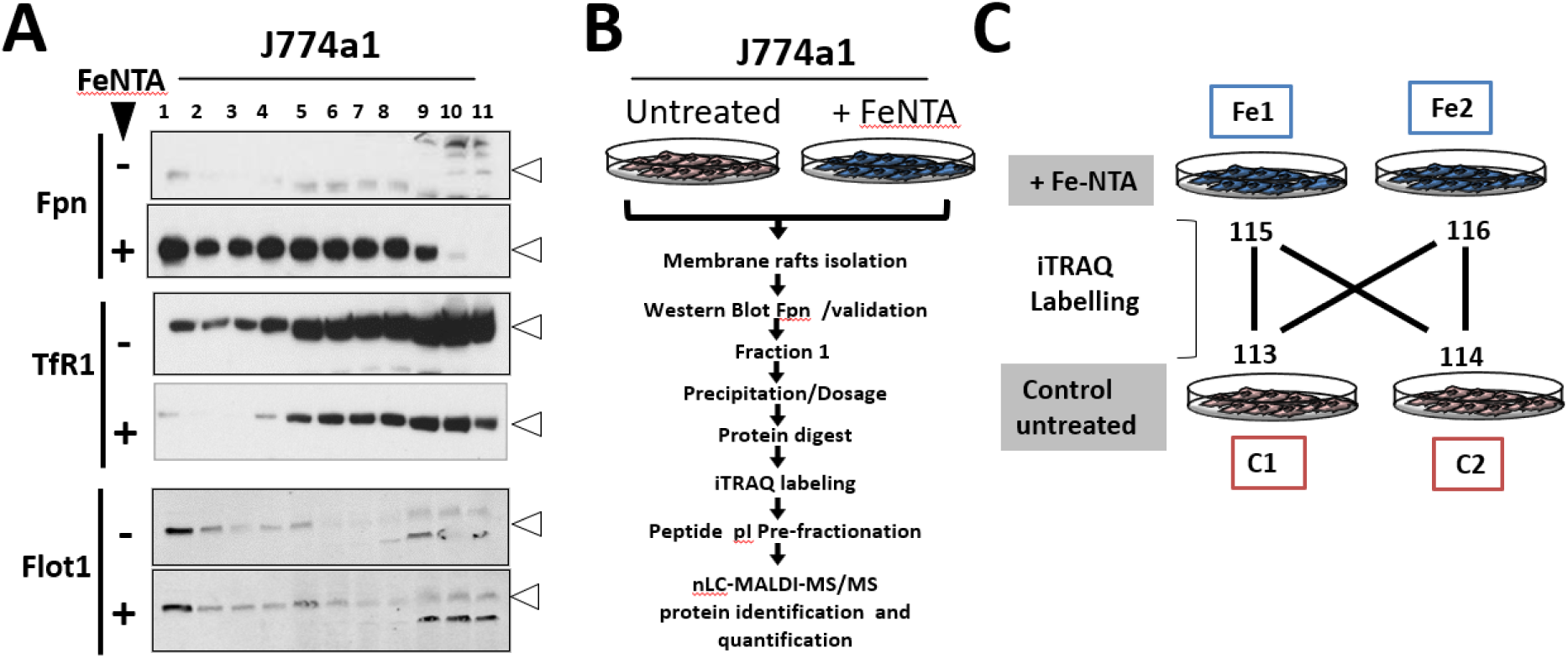
Experimental design for quantitative proteomic analysis of membrane rafts following iron loading in J774a1 macrophages. **(A)** Western blot analysis of J774a.1 macrophage cell fractions showing ferroportin (Fpn), transferrin receptor 1 (TfR1), and flotillin1 (Flot1) expression before (−) and after Fe–NTA (+) treatment in iodixanol gradient fractions (1-11). **(B)** Schematic representation of the treatment protocol for J774a.1 macrophage cells with FeNTA, followed by extraction/isolation of membrane rafts (lipid rafts defined as detergent-resistant membranes, DRM). Fraction 1 was collected and processed for the different steps of for proteomic analysis (see M&M). **(C)** iTRAQ labeling and data analysis. Each sample was prepared in duplicate (1 and 2). Each duplicate was labeled with a distinct iTRAQ isotope tag, allowing the calculation of four independent ratios to assess protein regulation between untreated (Control; C1, C2) and iron-treated (Iron; Fe1, Fe2) conditions: Ratio 1 = Fe1 / C1; Ratio 2 = Fe2 / C1; Ratio 3 = Fe1 / C2; Ratio 4 = Fe2 / C2 corresponding to tags 115:113, 116:113, 115:114, and 116:114, respectively. Each ratio was associated with a p-value and an error factor (EF; see Supplemental table 1).

#### Isobaric peptide labeling

The volume corresponding to 100 μg of protein from each sample was dried down in a centrifugal vacuum concentrator (Eppendorf). Samples were then resuspended in 25 μl of 500 mM TEAB (AB Sciex) containing 0.1% SDS. Proteins were then reduced, alkylated, digested, and labeled with iTRAQ reagents according to the manufacturer’s instructions (iTRAQ Reagents Application Kit, AB Sciex). The enzyme-to-substrate ratio was set to 1:10, and the pH was carefully controlled to ensure complete digestion. Each sample was labeled with a distinct isobaric tag; Control samples (C1, C2) and treated samples (Fe1, Fe2) were labeled with reporter ions 113, 114, 115, 116, respectively (Figure 1C). Labeled samples were subsequently pooled, and dried down in a vacuum concentrator prior to further analysis.

#### Peptide fractionation

Excess hydrolysed iTRAQ reagents were eliminated using a strong cation exchange (SCX) column (AB Sciex). Briefly, the SCX column was prewashed with 2 ml of cleaning buffer (25% Acetonitrile (Carlo Erba), 10 mM KH2PO4 (Carlo Erba), 1 M KCl, pH 3) and then equilibrated with 2 ml of loading buffer (25% Acetonitrile, 10 mM KH2PO4, pH 3). Sample was resuspended in 1 ml of loading buffer and acidified with few microliters of 10% H2PO4 to reach pH 3. Sample was then loaded onto the column and washed with 2 ml of loading buffer. Retained peptides were eluted with 500 µl of elution buffer (25% Acetonitrile, 10 mM KH2PO4, 350 mM KCl, pH 3). Eluted peptides were subsequently desalted using a Sep-Pak C18 cartridge (Waters). Briefly, the cartridge was activated with 3 ml of 90% acetonitrile containing 0.1% trifluoroacetic acid (TFA; Fluka) and equilibrated with 3 ml of 0.1% TFA. Peptides were loaded onto the cartridge, washed with 2mlof 0.1% TFA, and eluted with 1 ml of 70% acetonitrile, 0.1% TFA. The eluate was finally dried down using a vacuum concentrator (Eppendorf).

Peptides were fractionated by off-gel isoelectric focusing using 13 cm immobilized pH gradient (IPG) strips with a pH range of 3-10, according to the Agilent 3100 Off-Gel Fractionator quick-start guide. After focusing, fractions were collected individually. To recover peptides trapped within the IPG strip gel, 200 µl of extraction solution (50% methanol, 1% formic acid) were added to each well and incubated for 30 min at room temperature. The extracted peptides were then pooled with their corresponding fractions and dried down using a centrifugal vacuum concentrator.

#### Offline mass spectrometry analysis

Offline mass spectrometry analysis was performed by separating peptides using reverse-phase nano-liquid chromatography, followed by fractions collection, and subsequent analysis by tandem mass spectrometry (MS/MS) as describe below:

##### • Reverse-phase nano liquid chromatography fractionation

Each dried off-gel peptide fraction was resuspended in 100 µl of 0.1% TFA (Fluka) containing 10% acetonitrile (Carlo Erba). Ten percent of each sample was injected in duplicate onto an Ultimate 3000 nano-HPLC system (Dionex). Peptides were first concentrated and desalted on a C18 PepMap pre-column (0.3 mm I.D.× 5 mm, 100 Å pore size, 3 μm particle size; Dionex) at a flow rate of 30 μl/min using 0.1% TFA with 2% acetonitrile. They were then separated on a C18 PepMap 100 analytical reverse-phase column (75 μm I.D. × 150 mm, 100 Å pore size, 3 μM particle size; Dionex) at a flow rate of 300 nl/min. The mobile phases consisted of solvent A (0.1% TFA, 2% acetonitrile) and solvent B (20% solution A mixed vol/vol with 80% acetonitrile). After equilibration at 7% solvent B, a multi-step gradient was applied starting 3 min after injection: 16% B at 14 min, 19% at 22 min, 23% at 25 min, 32% at 51 min, 50% at 65 min, followed by a plateau at 95% B until 79 min. Fractionation was performed in duplicate for each off-gel peptide fraction using a Probot automated fraction collector (Dionex). Fraction collection started 18 min after injection and was performed directly onto a MALDI target plate. Fractions were collected every 10 s, resulting in a total of 384 spots per fraction. The LC eluent was mixed on-target with the matrix solution. The α-cyano-4-hydrocinnamic acid (CHCA) matrix solution was prepared at 2 mg/ml in 70% acetonitrile containing 0.1% TFA and 90 μM Glu-fibrinopeptide B for internal calibration (m/z = 1570.677).

##### • MS spectrum acquisition

Mass spectra were acquired using a 4800 MALDI-TOF-TOF mass spectrometer (AB Sciex). Data acquisition and processing were performed with the 4000 Series Explorer software (version 3.5.28193 build 1011; AB Sciex) in positive reflectron mode, using fixed laser fluence, low mass gate, and delayed extraction. External plate calibration was performed using four calibration points spotted across the plate, and additional internal calibration of spectra was achieved using the co-deposited Glu-fibrinopeptide. For each fraction, spectra were acquired in the mass range of 850 - 4000 Da at a laser shot frequency of 200Hz. Fifty spectra were acquired per step, with a total of 500 spectra summed per fraction. Spectra were processed to extract monoisotopic peak values from isotope clusters using a minimum signal to noise ratio of 20.

##### • Tandem MS and data-dependent tandem MS

For each MS spectrum generated from the nano-LC fractions, the eight most abundant precursor ions were selected for fragmentation, starting with the least abundant ions. precursors ions with a minimum signal-to-noise ratio threshold of 15 or within 200 resolution units of neighboring ions were excluded. For each selected precursor, 1000 MS/MS spectra were acquired in increments of 50 and summed. Spectra were processed to baseline subtraction and Savitsky–Golay smoothing (polynomial order 4 and 15 point across peak). To enhance the detection of lower abundant proteins, an exclusion list was applied during the duplicate fraction series. This list, generated from the first series of analyses, contained the m/z values and elution times of peptides that had been confidently identified and quantified.

##### • Protein identification and quantification

All acquisitions were combined and protein identification and quantification were carried out using the ProteinPilot software version 2.0.1 (Applied Biosytems, MDS-Sciex, Foster City, CA, USA). Database searches were initially performed against the Mouse International Protein Index (IPI) database (version 3.38) concatenated with a contaminant protein database. Since IPI identifiers are no longer maintained, all protein identifications were subsequently mapped to the corresponding current UniProt accession numbers, which are reported in supplemental Table 1. Data were processed using the following criteria: trypsin cleavage specificity, methylthio-modified cysteines, biological modifications enabled, and “thorough” search effort. Proteins were considered confidently identified if at least two unique peptides with >95% confidence were assigned and the overall protein confidence exceeded 95% (unused protein score>1.3). Differentially expressed proteins were defined as those with an iTRAQ-based fold change greater than 1.2 and the p-value was less than 0.05.

### Western blot

Protein extract (membrane, cytosol, DRM or biotinylated proteins) were solubilized in 1× Laemmli buffer and incubated for 30 min at room temperature prior to separation by SDS-PAGE and transfer onto PVDF membranes. DRM fractions were loaded in equal volumes per lane for SDS-PAGE analysis. For membrane, cytosol and biotinylated protein extracts, similar protein loading and transfer efficiency were assessed by Ponceau S staining or immunoblotting of actin or vinculin. PVDF membranes were blocked in TBST containing 7% non-fat dry milk and incubated overnight at 4°C with primary antibodies. After washing, membranes were incubated with HRP conjugated (chemiluminescence) or IRDye conjugated (Odyssey technology) secondary antibodies for 1 h at room temperature, and immunoreactive bands were detected using standard chemiluminescence methods or the Odyssey Infrared Imaging System. Antibodies and dilutions are listed in Supplemental Table 2.

### Immunofluorescence

Cells were cultured on 10 mm glass coverslips, washed three times with cold PBS, and fixed with 100% methanol for 15 min at −20 °C. After three PBS washes, cells were permeabilized or not with PBS containing 0.1% (in PBS) Triton X-100 for 10 min. Nonspecific binding sites were blocked for 45 min at room temperature with PBS containing 10% goat serum and 1% BSA. Cells were then incubated overnight at 4°C in a humidified chamber with one or more primary antibodies diluted in blocking buffer. After three washes with PBS containing 0.5% BSA, cells were incubated for 1 h at room temperature, protected from light, with secondary antibodies diluted in blocking buffer. Cells were finally washed three times with PBS/0.5% BSA, and coverslips were mounted onto glass slides using mounting medium. Primary and secondary antibodies and dilutions are listed in Supplemental Table 2.

### In cell western Blot (ICW; Odyssey)

BMDM cells were cultured in 96 well plates and treated as indicated. Cells were washed three times with cold PBS and fixed with 150 µl of 100% methanol per well for 15 min at −20°C. After three PBS washes, cells were permeabilized with PBS containing 0.1% Triton X-100 for 10 min at room temperature, followed by three PBS washes. Nonspecific binding was blocked with Odyssey Blocking Buffer (150 µl/well) for 90 min at room temperature. Cells were then incubated overnight at 4°C with primary antibodies diluted in blocking buffer. After five washes with PBS containing 0.5% Tween-20, cells were incubated for 1 h at room temperature, protected from light, with secondary antibodies diluted in blocking buffer supplemented with DRAQ5 (1:2000) for cell density normalization. After five final washes with PBS/0.5% Tween-20, plates were scanned using the Odyssey imaging system. Antibodies and dilutions are listed Supplemental Table 2.

### Use of Artificial Intelligence AI

AI (ChatGPT) was used for language polishing of this manuscript, while all content was originally created and reviewed by the authors. It also helped to transform a hand-drawn sketch into the graphical abstract.

## Results

### Isolation and Characterization of detergent resistant membrane fractions (DRM) containing the Iron exporter Fpn

To investigate the impact of iron overload on the macrophage membrane protein composition, the cell line J774a1 was treated or not (control) with iron (FeNTA) and lipid raft as detergent resistant membrane (DRM) from these cells were isolated using a discontinuous iodixanol gradient ultracentrifugation (Fig.1). DRM are low density membrane fractions that remain insoluble after extraction with non-ionic detergent (Triton X-100) at +4°C and are isolated by density-gradient ultracentrifugation. Despite not totally equivalent to native lipid rafts, DRM are a biochemical readout of membrane microdomains enriched in lipids raft proteins (Schuck et al., 2003). Eleven DRM fractions were collected from the top to the bottom of the gradient, corresponding to increasing density (fractions 1-11).

Western blot analysis of the gradient fractions from J774a1 (Fig.1A) revealed that, as expected, the known raft marker flotillin1 was enriched in lower-density fractions (mainly fraction 1) in both control and iron overload condition. On the other hand, the transferrin receptor 1 (TfR1), a non-raft marker, was strongly enriched in high-density non-DRM fractions (fractions 9 to 10 notably). These distributions were consistent with effective membrane fractionation. As expected, TfR1 expression was markedly reduced upon iron treatment, consistent with its well-established negative regulation mediated by the IRP-IRE system (Muckenthaler et al., 2008). Importantly, as previously described (Auriac et al., 2010), Fpn was barely detectable under basal conditions but it was strongly upregulated by FeNTA, with the induced protein enriched in low-density iodixanol fractions (1-4). A longer exposure of the blot indicated the presence of Fpn in DRM and non-DRM (NDRM) fraction in basal condition in J774a1 (Fig.S1).

Given the strong enrichment of Fpn in the lightest iodixanol fraction 1, this fraction was selected for downstream proteomic analysis. Lipid raft fraction 1 isolated from control and FeNTA treated J774a1 macrophages were then analyzed by iTRAQ-based quantitative proteomics coupled to nLC-MALDI-MS/MS, allowing comparison of protein abundance across biological duplicates using multiple independent ratios (Fig.1B & C).

### Quantitative proteomic analysis in J774a1 treated or not with iron

As shown in Fig.2A, iTRAQ labeling efficiency reached 99.6% and we identified 1013 and 823 proteins with greater than 95% and 99% confidences, respectively. The complete list of detected protein is given in Supplemental table 1. In Fig.2B, all proteins were then ranked according to their fold change between iron-treated and control conditions (ratios with lowest p-value), and the resulting distribution was visualized as a ranked bar plot of protein fold changes. This global profile indicates an important number of proteins exhibiting negative as well as positive regulation. Indeed, 136 identified proteins were positively upregulated including 79, 66 and 21 proteins with a ratio of expression superior to 2, 5 and 20, respectively (Fig.2B).

**Figure 2.**
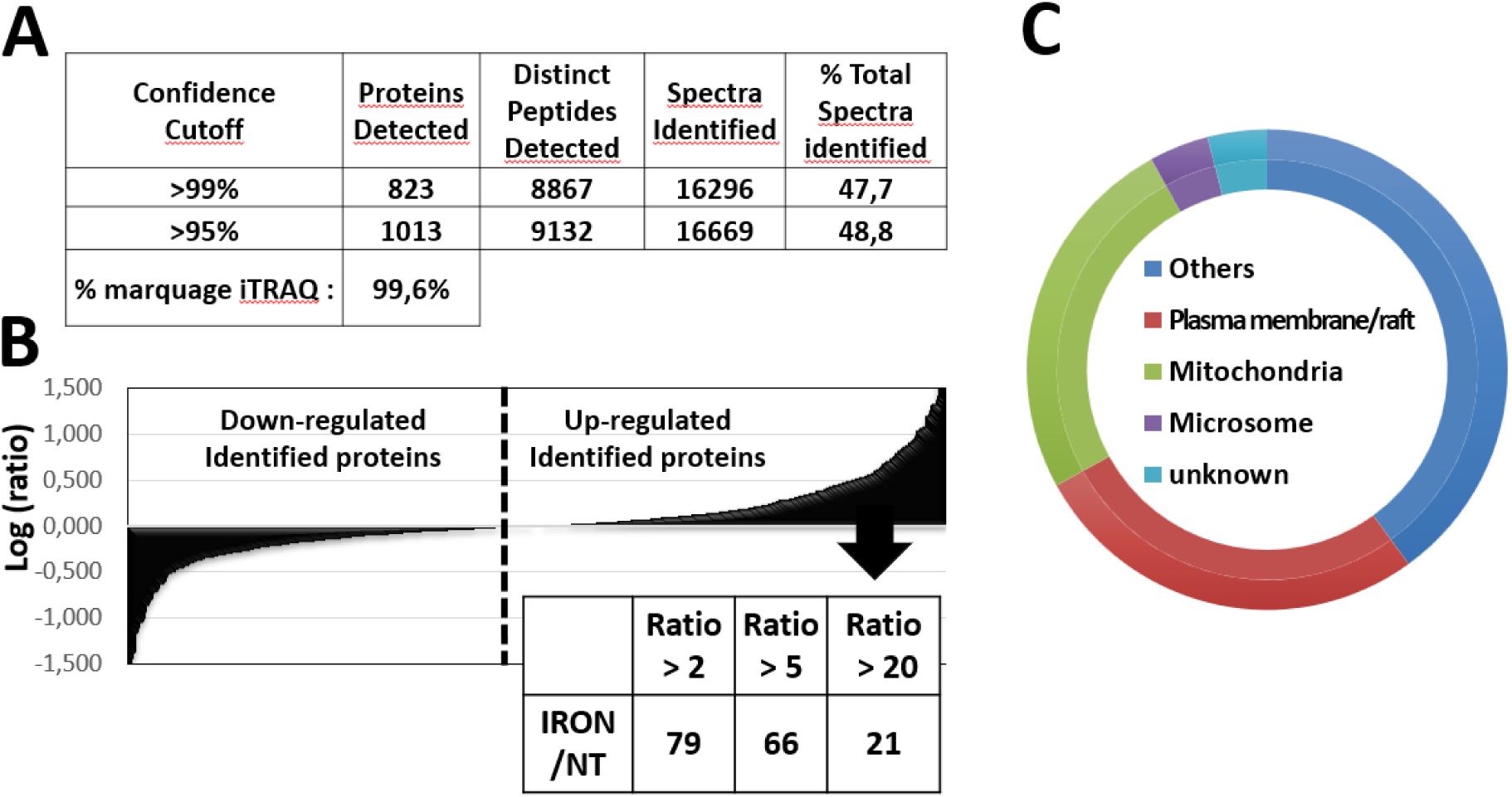
Protein identification, quantification, and subcellular distribution under iron versus control (UT; untreaded) conditions in J774a1 cells. **(A)** *Protein identification and quantification:* The table summarizes parameters of protein detection under two confidence thresholds (>99% and >95%), including the total number of proteins identified (823 and 1,039, respectively), distinct peptides, spectra identified, and the percentage of total spectra matched. The iTRAQ labeling efficiency, indicating labeling completeness, is 99.6%. **(B)** *Ranked fold-change (waterfall) plot:* The bar chart depicts the logarithmic fold changes in protein abundance (log2 scale) in the iron-treated condition relative to control (UT). Negative values indicate downregulated proteins, while positive values indicate upregulated proteins in response to iron exposure. The accompanying table (below) indicate number of proteins exhibiting abundance ratios greater than 2, 5, and 20, illustrating differential upregulation magnitudes. **(C)** *Subcellular distribution of identified proteins:* The doughnut chart presents the distribution of identified proteins classified by subcellular localization or functional compartments: plasma membrane/raft (red), mitochondria (green), microsomes (purple), unknown (cyan), and others (blue). This distribution highlights the cellular compartments predominantly affected or represented in the dataset under the iron treatment.

We then performed a classification analysis of the identified proteins based on their annotated subcellular localization and functional compartment (Fig.2C). The resulting chart illustrated the distribution of proteins identified following iron treatment. A substantial proportion of proteins was associated with the plasma membrane and lipid raft fraction whereas mitochondrial and microsomal proteins constitute a notable fraction. A smaller subset of proteins could not be assigned to a defined compartment and was classified as unknown, while the remaining proteins were into other cellular localizations.

A list of the 79 proteins upregulated (≥2-fold) in response to iron treatment is shown in Supplemental table 3 and in Fig.3A. Protein ranking and selection in Supplemental table 3 & Fig.3 were based on their positive ratio associated with the lowest p-value (< 0.05). In Fig.3B, the bar graph displays a focus of a subset of 60 upregulated proteins with fold increases below 35, ranked by fold induction and labeled by gene name. Proteins exhibiting fold changes greater than 35 were excluded from the quantitative analysis. Such extreme ratios are difficult to interpret reliably in iTRAQ-based proteomics, as they can result from very low reporter-ion intensities in one of the compared conditions, leading to unstable and disproportionately large ratios. These extreme ratios may therefore reflect limitations in reporter-ion quantification and signal-to-noise rather than genuine biological differences. To minimize the risk of overinterpreting potentially unreliable quantitative measurements, proteins with fold change >32 were consequently not considered for downstream analyses. Importantly and as a control, we detected our protein of interest Fpn in our raft fraction and observed a strong increased expression after iron treatment (25x). Both flotillin 1 and 2 were also detected in our analysis, confirming the raft nature of our samples. Further protein distribution such as peroxiredoxin-1 (Prdx1) and glucose-6-phosphate dehydrogenase (G6pdx/G6PD) suggest a stress and metabolic response. Interestingly, another protein involved in cellular stress and in iron metabolism as well, the heme oxygenase 1 (Hmox1) was clearly identified in our analysis. Indeed, parallel to Fpn, Hmox1 was strongly upregulated (up to 20-fold change). Intriguing, the ferritin heavy chain (FtH), a well know cytosolic protein, was also detected in the lipid raft fraction 1 analyzed and we observed a marked increase following iron treatment. However, FtH enrichment was significantly observed only in two out of the four quantitative conditions analyzed (Supplemental table 1).

**Figure 3.**
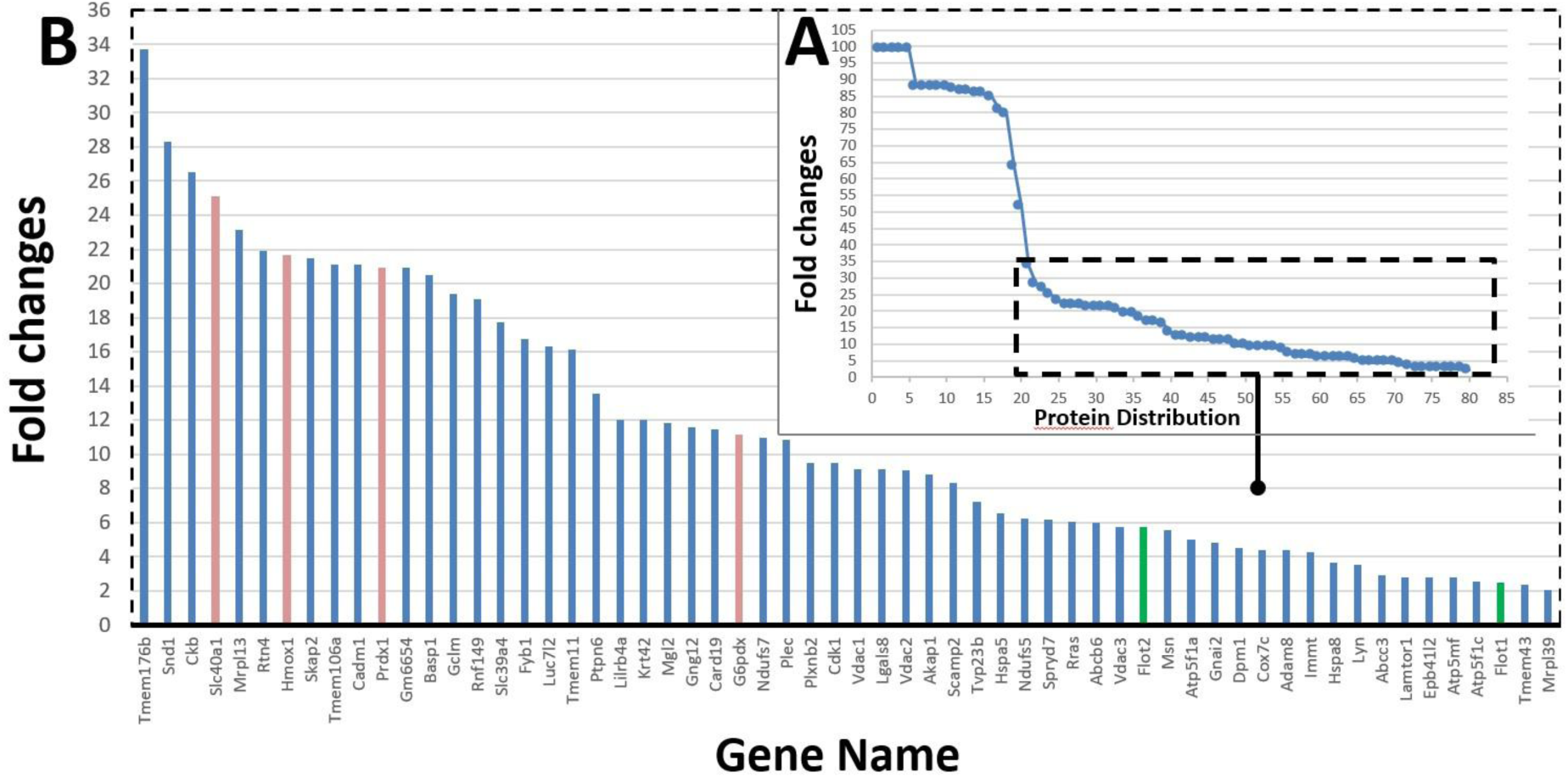
Histogram Representation of the 79 proteins exhibiting more than a twofold increase. **(A)** The graph in the upper right shows the global distribution of all proteins exhibiting more than a twofold increase (79 proteins in total) identified by iTRAQ-based proteomic analysis (see supplemental Table 3, sheet Figure 3). **(B)** The larger bar graph below displays a focus (box in A.) of a subset of 60 upregulated proteins with fold increases below 34, ranked by fold induction and labeled by gene name. Bars in orange color indicate proteins of particular interest discussed in the text. Bar in green correspond to lipid raft marker Flotillin 1 & 2. Ratios of each protein correspond to the one obtained with the lowest p-values among the four calculated ratios.

We next increased the stringency of the analysis by ranking proteins that exhibited a positive fold change across all four calculated ratios (Fig.1C) and that were associated with four, three, or two statistically significant p-values (Fig. 4). Under these criteria, Hmox1, Prdx1 and G6pdx showed increased abundance in all four ratios, with each ratio reaching statistical significance (p < 0.05). Fpn also displayed increased abundance in all four ratios, but only two of these reached statistical significance.

**Figure 4.**
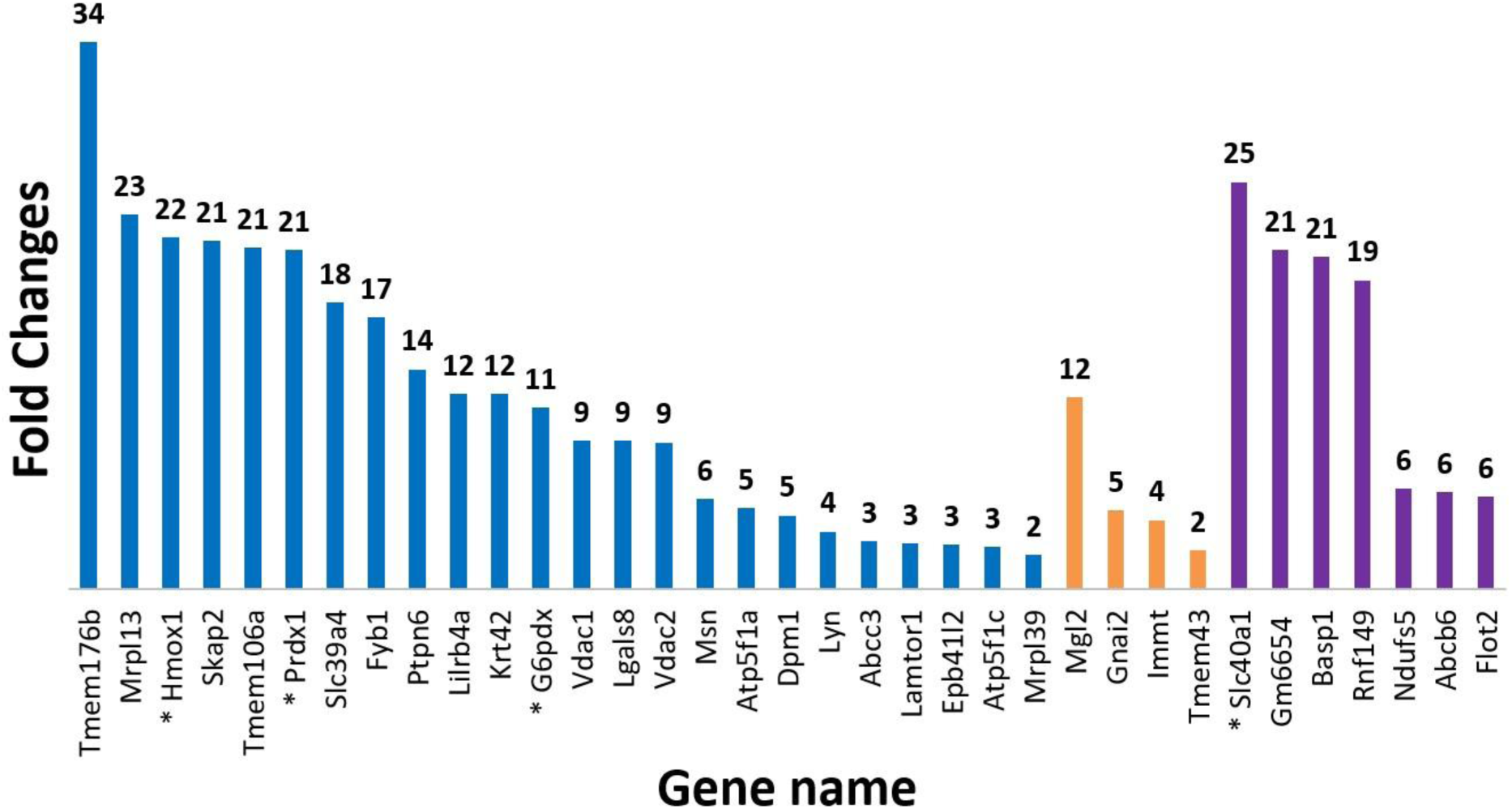
Identification of proteins upregulated under iron treatment in the four calculated ratios. Bar graph showing 35 proteins identified by iTRAQ-based proteomic analysis with fold changes greater than or equal to 2 and up to 34 in the four calculated ratios (see Supplemental table 3, sheet figure 4). For each protein, the ratio with the lowest p-value among the four measurements was selected. Blue bars indicate proteins whose increase was observed in all four analyses with statistically significant p-values in each case. Orange bars represent proteins increased in all four conditions but with significant p-values in only three conditions. Purple bars correspond to proteins increased in all four conditions but reaching statistical significance in only two. * Indicates protein of interest.

### Biochemical and cellular evidences of Hmox1 expression increase with iron overload

Due to its role in heme degradation and iron release, Hmox1 was of particular interest, and subsequent analyses focused on its further characterization. We first confirmed the upregulation of Hmox1 in our experiment condition in macrophages. For such purpose, we used bone marrow derived cells (BMDM) from DBA2 and SWISS mice and we treated them or not (control) overnight with iron-NTA (Fig.5). Western blot analysis (Odyssey) illustrated a strong increase of Hmox1 in response to iron. Such increase was confirmed by In cell western blotting (Fig.5B & C) as well as by immunofluorescence (Fig.5D). Hmox1 induction was concomitant with induction of Fpn (Fig.5C) and ferritin (Ft, Fig.S2). ICW demonstrated a clear dose-response induction of Fpn, FtH and Hmox1 protein levels when macrophages were treated with increasing FeNTA concentrations (0, 25, 50, 100 µM) (Fig.5C and Fig.S2).

**Figure 5.**
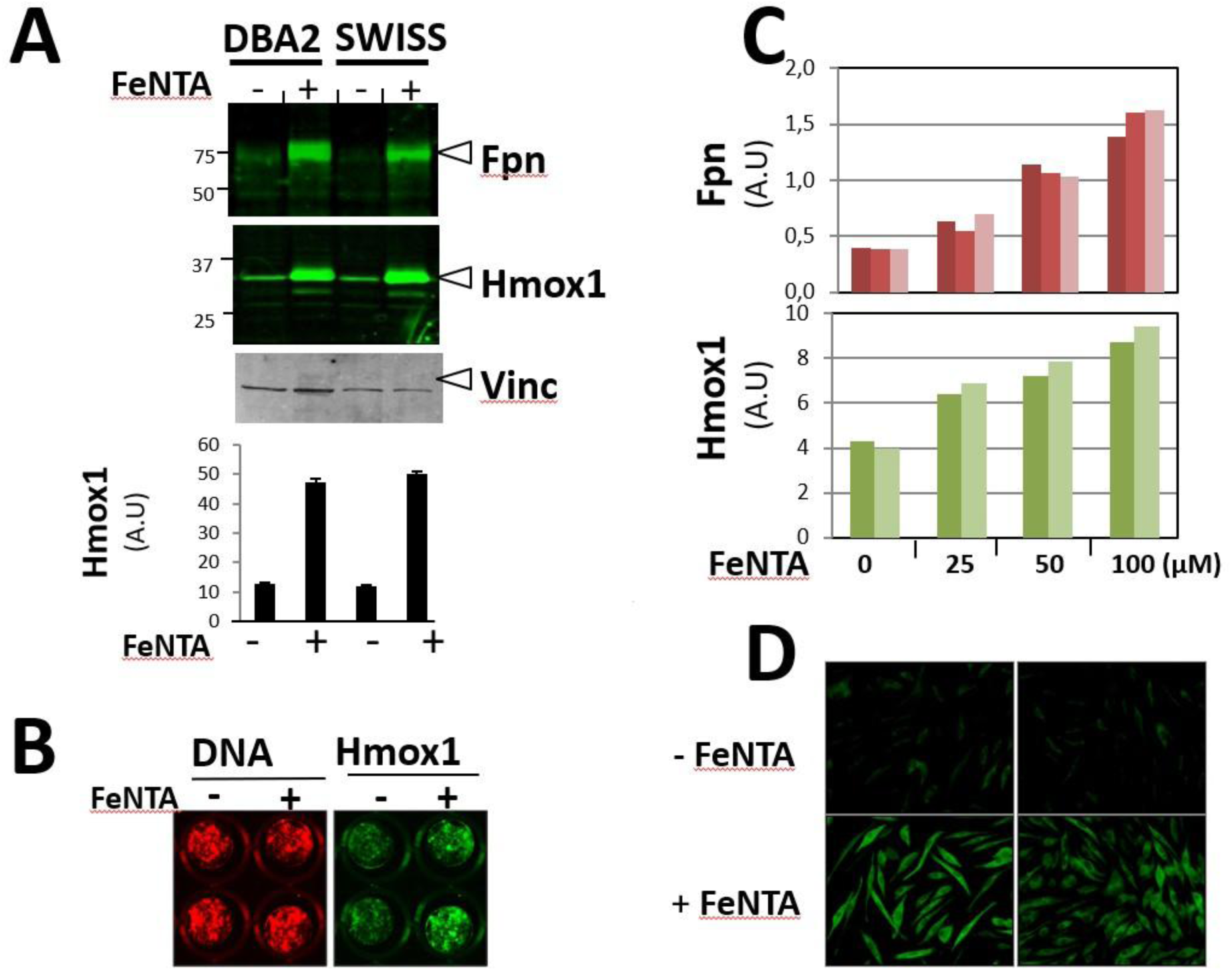
Effect of FeNTA exposure on ferroportin (Fpn) and heme oxygenase-1 (Hmox1) expression in two mouse strains. **(A)** Representative western blot analysis of Fpn, Hmox1, and vinculin (loading control) in DBA2 and SWISS in membrane protein extract from bone marrow derived macrophages BMDM treated (+) or not (-) with FeNTA. Vinculin is used as a loading control. Densitometric quantification of Hmox1 expression is shown below (normalized to vinculin expression; arbitrary units, A.U.). **(B)** Representative image of “In cell western blot” (Odyssey) showing immunofluorescence staining of DNA (red, Draq5) and Hmox1 (green) in BMDM cells treated without (–) or with (+) FeNTA (2 wells for each condition are shown). **(C)** Fluorescence quantification (A.U.; normalization to DNA staining) from an In-Cell western Blotting showing a dose-dependent effects of FeNTA (0–100 μM) on Fpn (top panel, red bars, 3 independent wells) and Hmox1 (bottom panel, green bars, 2 independent wells) expression levels. **(D)** Classical Immunofluorescence detection of Hmox1 in BMDM cells cultured in the absence (–) or presence (+) of FeNTA. Images are representative of three independent experiments.

All together, these results confirm our proteomic findings and demonstrate that iron treatment greatly enhances expression of both Fpn and Hmox1.

### The transcription factor Nrf2 is a key player in the regulation of Fpn and Hmox1 by iron

Ingenuity Pathway Analysis was performed on the proteomic dataset (Fig.S3). This analysis indicated that the upregulated proteins Prdx1, Fpn, G6pd, Gclm, and Hmox1 detected in our experimental conditions are connected within a network centered on the redox-sensitive transcription factor Nrf2. Consistent with this network organization, several of these proteins are known targets of Nrf2-regulated pathways. Based on these observations, the involvement of Nrf2 in the regulation of Hmox1 and Fpn proteins was further investigated (Fig.6).

**Figure 6.**
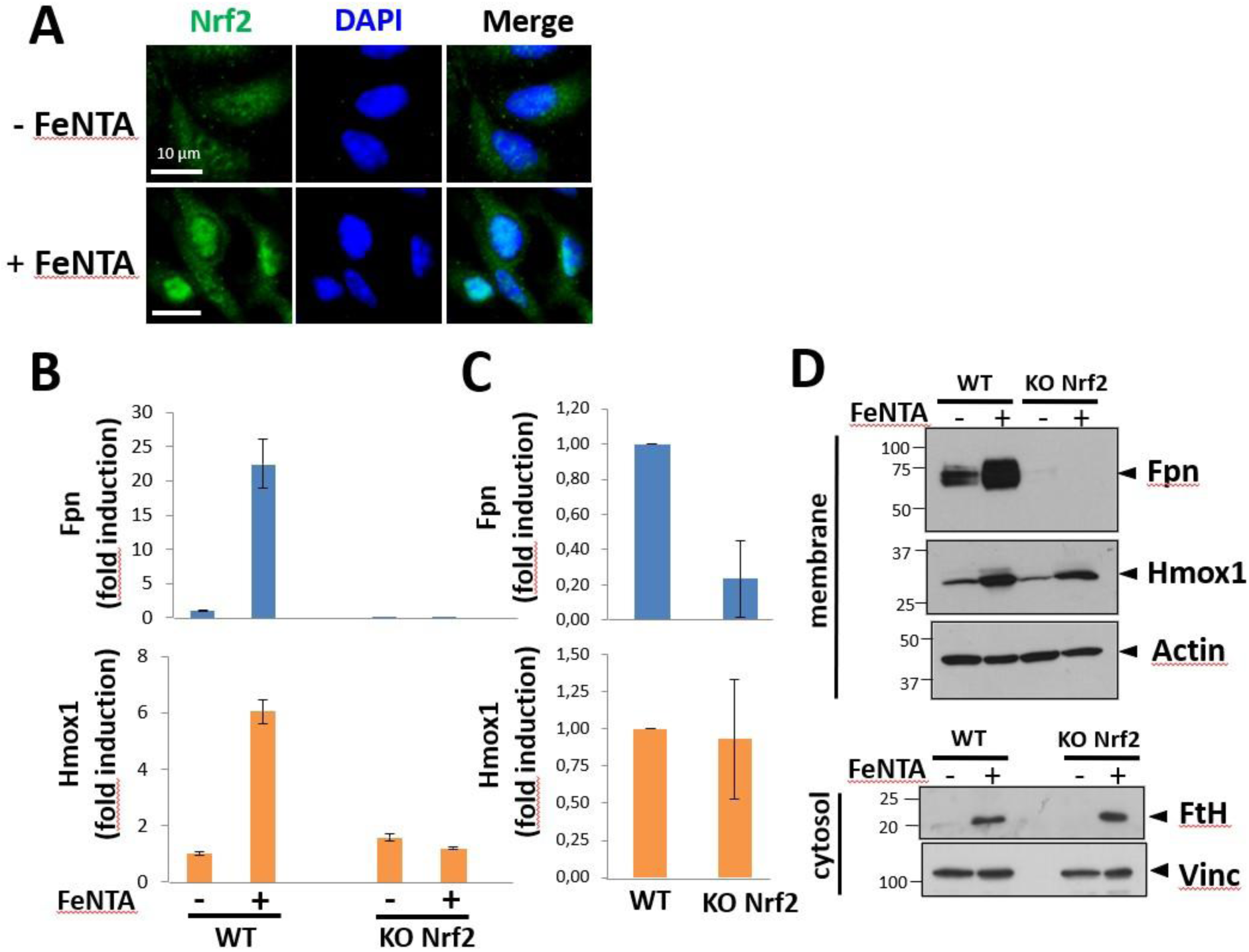
Impact of Nrf2 on the expression of ferroportin and heme oxygenase-1 in macrophages treated with iron. **(A)** Representative confocal images of macrophages stained for Nrf2 (green) and nuclei counterstained with DAPI (blue). Panels show control (untreated) and FeNTA-treated BMDM, with individual channels for Nrf2 and DAPI, as well as merged images. **(B)** Histograms showing mRNA expression levels of Ferroportin (Fpn) and Heme Oxygenase-1 (Hmox1) measured by quantitative PCR (qPCR) in bone marrow-derived macrophages (BMDMs) from wild-type (WT) and Nrf2 knockout (KO) mice. Cells were either untreated or treated with FeNTA (iron) or heme. **(C)** The histograms represent a focused comparison of basal (untreated) mRNA expression levels between WT and Nrf2 KO BMDMs. **(D)** Western blot analysis of membrane and cytosolic fractions from wild-type (WT) and Nrf2 knockout (KO) macrophages. Upper panels show membrane extracts probed for Ferroportin (Fpn) and Heme Oxygenase-1 (Hmox1), with actin as a loading control. Lower panels show cytosolic extracts analyzed for Ferritin Heavy Chain (FtH) and vinculin (Vinc) as a loading control.

Immunofluorescence microscopy showed an increased accumulation of Nrf2 in the nucleus of macrophages following Fe–NTA exposure (Fig.6A). To further assess the contribution of Nrf2, mRNA (Fig. 6B.C) and protein levels of Fpn and Hmox1 (Fig.6D) were analyzed in wild-type and Nrf2 knockout (KO) BMDM after iron treatment. In wild-type cells, iron–NTA induced the expression of both Fpn and Hmox1 at the mRNA level, whereas this induction was completely abolished in Nrf2 KO macrophages. Under basal conditions, the Fpn mRNA expression was reduced in Nrf2-deficient cells, while the *Hmox1* mRNA basal expression was not affected. Western blot analysis of membrane proteins confirmed that the iron-induced upregulation of Fpn observed in wild-type macrophages was absent in Nrf2 KO macrophages (Fig.6D). As a control, ferritin, only detected in the cytosolic fraction, was increased following iron treatment in both wild-type and Nrf2 knockout macrophages, reflecting elevated intracellular iron levels in both conditions.

### Heme Oxygenase-1 is localized in detergent resistant membrane (DRM) in iron or heme treated macrophages

The detection of Hmox1 in our proteomic analyses strongly suggested the presence of the enzyme in a lipid raft compartment after iron treatment. Therefore, western blot analyses of iodixanol gradient fractions were then performed to examine the distribution of Hmox1 in DRM from both J774a1 macrophages (Fig.7A) and BMDM (Fig.7B). To facilitate protein detection and comparison, iodixanol fractions were pooled into DRM1 (fractions 1-4), DRM2 (fractions 5-8), and non-DRM (NDRM; fractions 9-11). This fractionation confirmed that Fpn was detected only under iron-treated conditions and was predominantly enriched in the DRM1 and DRM2 fractions, as confirmed by the presence of the raft markers caveolin1 in BMDM and flotillin1 in J774a1 cells (Fig.7). In both macrophage cultures, TfR1 was poorly or not detected in the DRM1 fraction but was present mainly in DRM2 and NDRM. Under basal conditions, Hmox1 was detected exclusively in the NDRM fractions in both J774a1 and BMDM. Upon iron treatment, Hmox1 expression was strongly induced and detected in DRM1, DRM2 and NDRM fractions in the two macrophage cultures. Figure.S4A presents western blot analyses of Hmox1 of all individual iodixanol gradient fractions from BMDM. Following iron treatment, Hmox1 was detected across all fractions of the gradient. Under basal conditions, Hmox1 was mainly detected in fractions 9 to 11, corresponding to the NDRM pool. A longer exposure of the western blot detection of Fpn in figure 7B indicated that the iron exporter is mostly present at low basal levels in DRM1 and DRM2, while a weak signal is observed in the NDRM pool (Fig.S4B).

**Figure 7.**
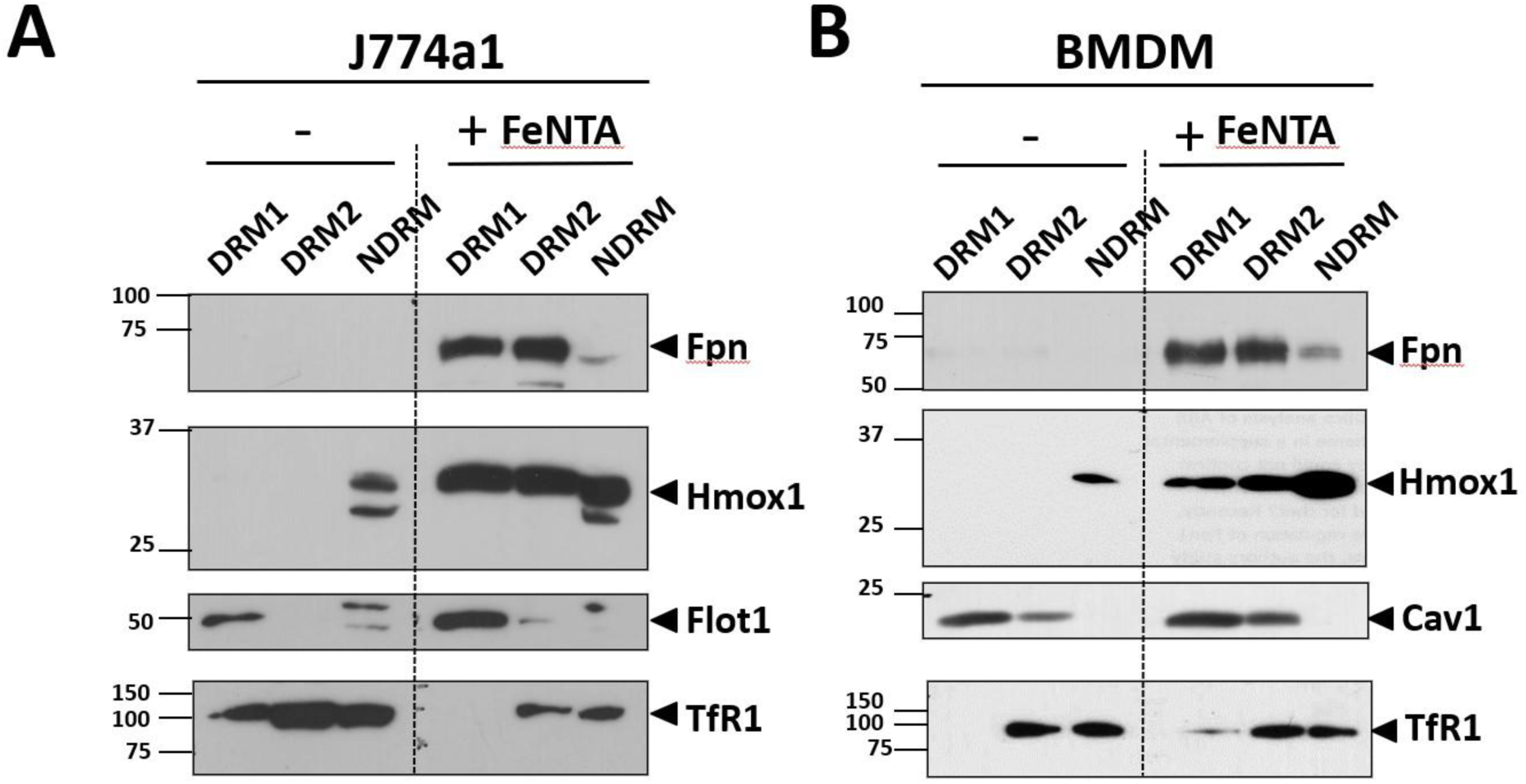
Distribution of ferroportin and Heme oxygenase proteins in detergent-resistant and non-detergent-resistant membrane fractions from J774a.1 and BMDM cells following FeNTA treatment. **(A)** J774a.1 macrophages and **(B)** bone marrow-derived macrophages (BMDM) were untreated (–) or exposed to FeNTA (+). Membrane fractions obtained by iodixanol gradient centrifugation were pooled into detergent-resistant membranes (DRM1, DRM2) and non-detergent-resistant membranes (NDRM). Equal volumes from each pooled fraction were analyzed by western blot for ferroportin (Fpn), heme oxygenase-1 (Hmox1), flotillin1 (Flot1, lipid raft marker for J774a.1) or caveolin1 (Cav-1, lipid raft marker for BMDM), and transferrin receptor 1 (TfR1, non-raft marker). Data are representative of at least two independent experiments for each macrophage population. Vertical dashed lines indicate the virtual separation of untreated and treated samples. Molecular weights are shown on the right.

To assess whether similar detections could be observed using a more physiological iron source, BMDM were treated or not with heme (overnight), and proteins were separated by iodixanol gradient fractionation (Fig.8). Western blot analysis of the 11 collected fractions showed that caveolin1 and TfR1 displayed distributions comparable to those observed with iron–NTA treatment, defining the same DRM1, DRM2, and NDRM pools. As previously observed, Fpn was not detected in untreated cells in such experimental condition, confirming its low basal expression level in BMDM. Following heme treatment, Fpn was strongly detected in DRM1, with a pronounced signal in fraction 1. In untreated macrophages, Hmox1 was detected only in the NDRM fractions as previously observed (Fig.7). Upon heme exposure, Hmox1 expression was strongly induced and detected across all gradient fractions, with detectable signal in DRM1, overlapping with Fpn expression.

**Figure 8.**
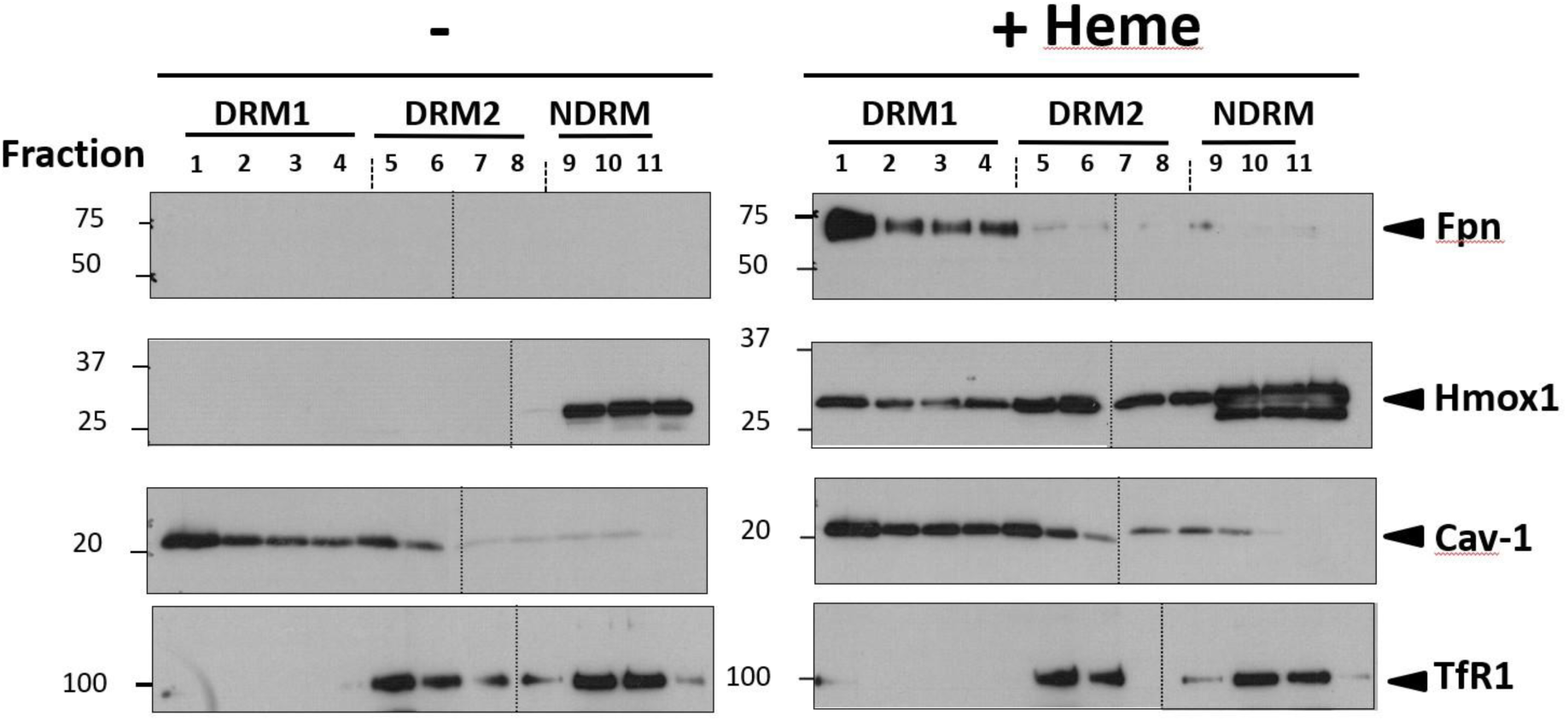
Distribution of Fpn and Hmox1 in DRM and NDRM membrane fractions from BMDM cells treated with heme. Cells were untreated (–) or treated with heme (+) before membrane fractionation by iodixanol gradient centrifugation. Eleven gradient fractions were collected and pooled into detergent-resistant membrane fractions (DRM1: fractions 1–4; DRM2: fractions 5–7) and non-detergent-resistant membranes (NDRM: fractions 8–11). Equal protein amounts from each fraction were analyzed by western blot for ferroportin (Fpn), heme oxygenase-1 (Hmox1), caveolin1 (Cav-1, lipid raft marker), and transferrin receptor 1 (TfR1, non-raft marker). Data are representative of at least two independent experiments for each protein. Vertical dashed lines indicate repositioned gel lanes. Molecular weights are shown on the right.

### Colocalization of Hmox1 and Fpn at the cell surface of iron and heme treated BMDM

Fpn was shown to be strongly enriched at the cell surface of macrophages after iron treatment (Delaby, et al., 2005; Knutson et al., 2003). Therefore, the presence of Hmox1 at the plasma membrane of macrophages following iron treatment was further examined using cell-surface biotinylation assays (Fig.S5). Biotinylation experiments revealed the presence of two Hmox1 protein species in the intracellular pool. In contrast, a strong signal corresponding to the upper form of Hmox1 was detected in the biotinylated (plasma membrane) fraction, together with Fpn, strongly present in this cellular compartment. As a control, FtH was detected exclusively in the intracellular fraction and was absent from the biotinylated fraction.

Since Hmox1 and Fpn were detected within the same cellular compartments (DRM and cell surface), their potential co-localization was examined in BMDM treated with either iron or heme using confocal microscopy (Fig.9). We first analyzed the co-distribution of Hmox1 or Fpn with the lipid raft marker caveolin1 (Fig.9A and Fig.S6) in BMDM. Caveolin1 displayed a punctate staining pattern, whereas Hmox1 showed a more granular intracellular distribution (Fig.9A). Nonetheless, partial overlap between Hmox1 or Fpn with caveolin1 signals was observed. This overlap was illustrated by line-scan pixel intensity profiles acquired along defined regions within a single optical plane, showing coincident signal peaks for both proteins (see Materials and Methods). No co-localization between Hmox1 and caveolin1 was detected in untreated cells (Fig.S6). Co-localization was observed only following iron or heme treatment, concomitant with the induction of Hmox1 expression (Fig.S6).

**Figure 9.**
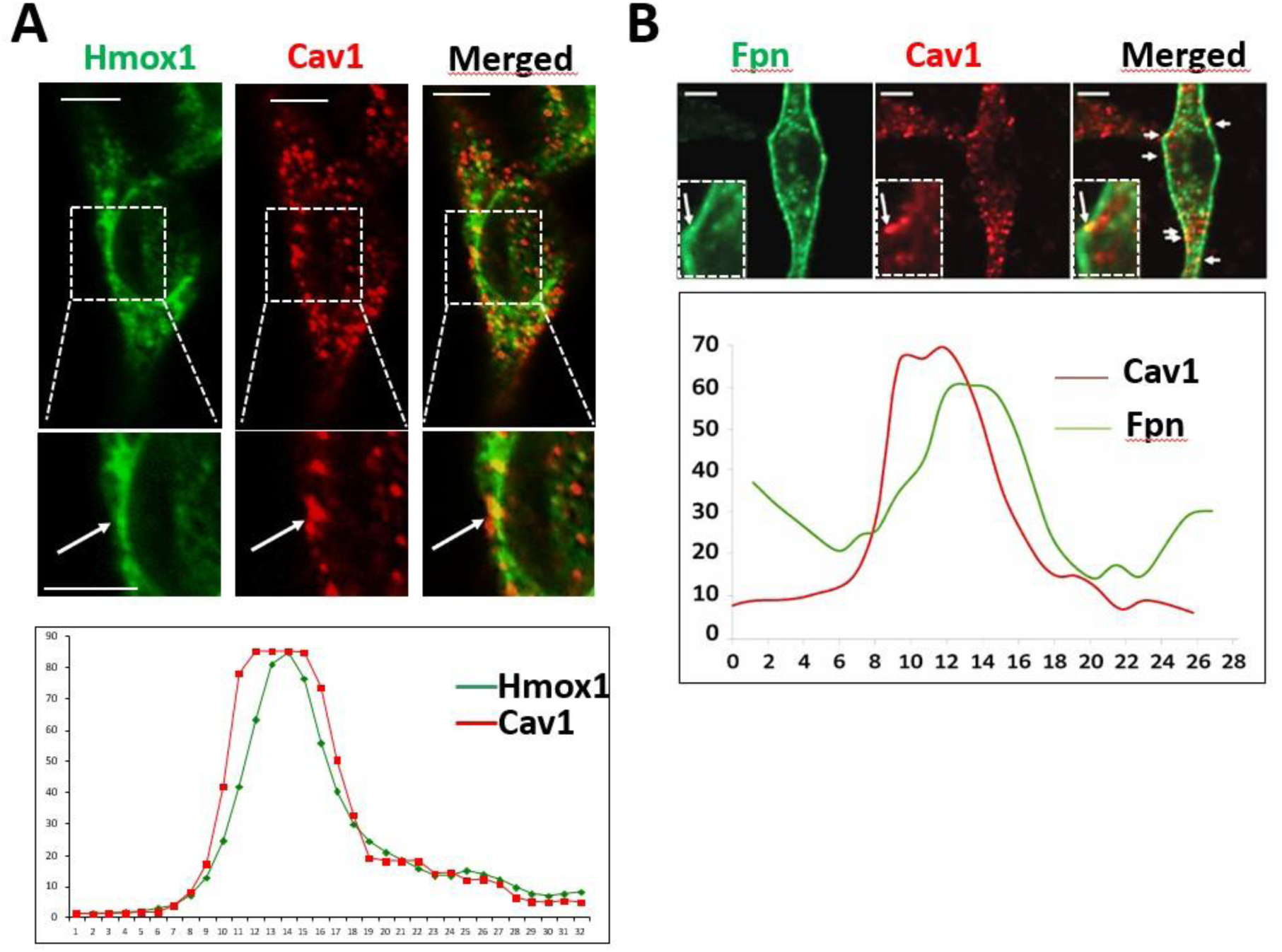
Partial colocalization of heme oxygenase-1 with ferroportin and caveolin1 in macrophages analyzed by confocal microscopy. Confocal imaging of heme oxygenase-1 (Hmox1), ferroportin (Fpn) and caveolin1 (Cav1) in BMDM treated either with FeNTA. White arrows indicate regions of partial colocalization between Hmox1 or Fpn with caveolin1, suggesting spatial proximity or potential interaction of these proteins under these treatment conditions. The graph in (A) shows a pixel intensity profiles across a specific focal plane illustrating the colocalization of Hmox1 with caveolin1. In (B), the pixel intensity profiles confirm the colocalization of Fpn with Caveolin1 after iron treatment. Bar = 5 µm

Interestingly, Z-stack analyses (Fig.10) revealed compartments positive for both Fpn and Hmox1 (yellow staining), predominantly located at the cell surface following FeNTA or heme treatment (Fig.10A). Under these conditions, Fpn staining was mainly observed at the plasma membrane, whereas Hmox1 was distributed throughout the cell but with detectable overlap in discrete specific plasma membrane regions (arrow heads). Additional representative confocal images with line-scan pixel intensity profiles showing Fpn and Hmox1 colocalization at the cell surface of BMDM treated with FeNTA are presented in Fig.10B.

**Figure 10.**
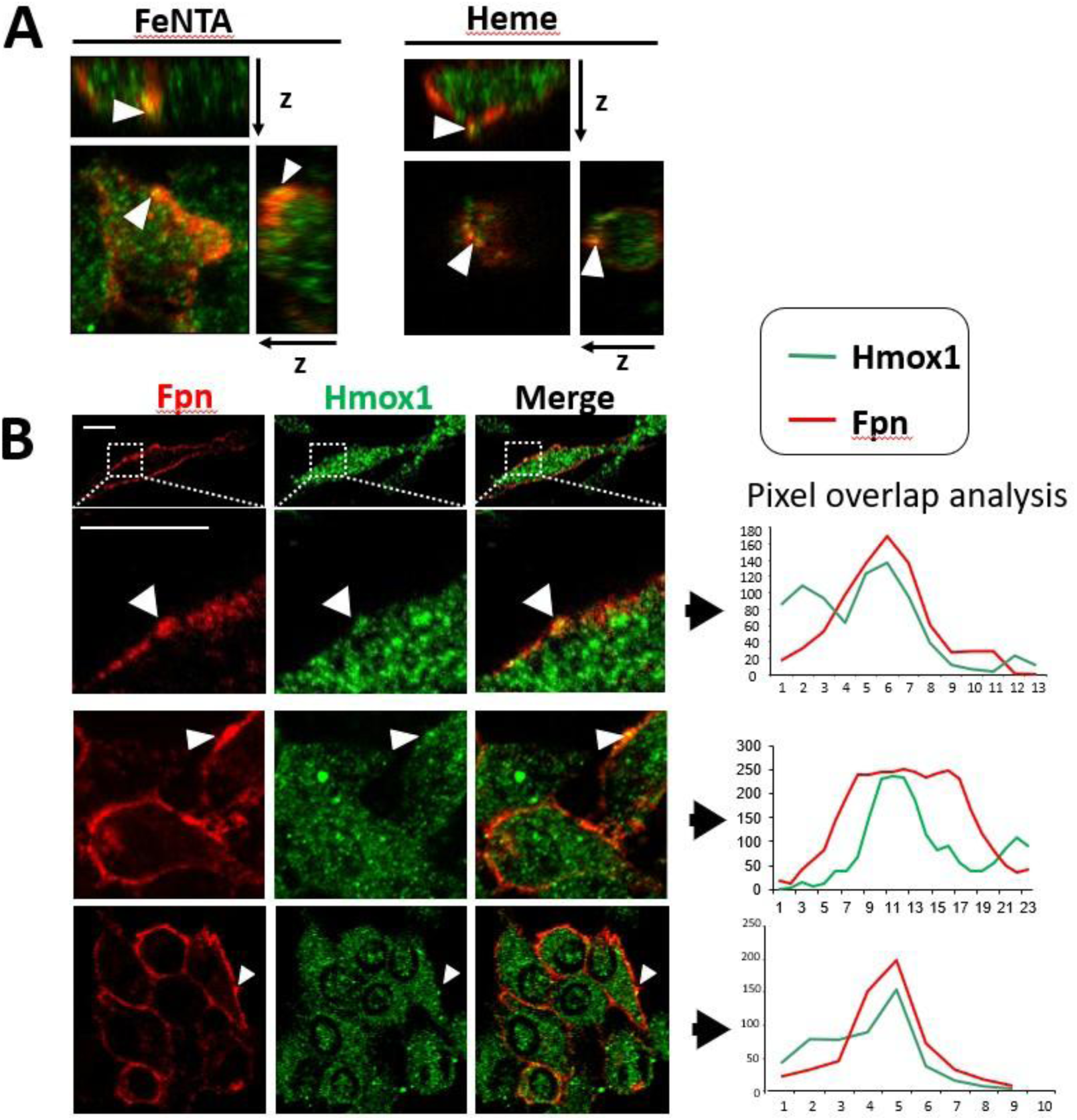
Colocalization analysis of Fpn and Hmox1 by confocal microscopy. Representative confocal images showing **Fpn** (red) and **Hmox1** (green) in cells under FeNTA. Merged images (yellow/orange) reveal areas of co-localization (arrowheads). **(A)** Z-stack analysis of Hmox1 and Fpn co-localization. **(B)** Line-scan intensity profiles (right panels) were obtained along the indicated regions of interest (Arrowheads), illustrating the degree of overlap between **Fpn** and **Hmox1.** Bar = 5 µm

## Discussion

### An Iron induced Lipid raft proteome in macrophage

Membrane rafts are of particular interest as they concentrate specific plasma membrane proteins and lipids into defined microdomains that are involved in numerous cellular biological activities and cell signaling (Lingwood & Simons, 2010; Simons & Sampaio, 2011). Quantitative proteomics has been described as a powerful approach to the understanding of the biology of membrane microdomains (Zheng & Foster, 2009). Therefore, proteomic analyses of lipid rafts/DRMs in macrophages have been performed in various contexts, but not specifically in the context of iron treatment (Dhungana et al., 2009; N. Li et al., 2003). On the other hand, several global proteomic studies have been performed to assess the effects of iron on macrophage expression profiles, including the study of Polati et al. (2012), and more recently, the quantitative proteomics and phosphoproteomics of human macrophages treated with iron carbohydrate complexes (Bossart et al., 2023). However, to date, no dedicated proteomic analysis of the lipid raft/DRM fraction has been reported in iron-treated macrophages.

The expression of the iron exporter Fpn within macrophage membrane microdomains (Auriac et al., 2010) prompted us to investigate the quantitative (iTRAQ) proteomic composition of DRM containing Fpn. Because iron treatment markedly increases the presence of Fpn at the cell surface, we took advantage of this experimental condition to identify potential functional partners of Fpn within these membrane compartments. Our quantitative proteomic analyses were performed using the DRM samples isolated from J774a.1 cell line to take advantage of its clonal homogeneity, an important consideration for highly sensitive quantitative proteomic approaches. In contrast, BMDM may exhibit greater variability due to inter-animal differences and variations associated with cell differentiation in the presence of M-CSF. In J774a1 cells, the raft marker flotillin1 was enriched in the lightest fraction (fraction 1) where Fpn was also strongly detected. TfR1, used as a non-raft marker, was predominantly detected in the heaviest fractions (fraction 5 to 11). However, a weak TfR1 signal was occasionally observed in fraction 1. Thus, the distribution of TfR1 did not strictly correspond to a non-raft distribution. This apparent overlap should be interpreted in the context of density-gradient fractionation, which reflects a continuum of protein distribution rather than a separation into strictly discrete membrane compartments. Signal intensity may also be influenced by differences in the amount of protein loaded on the gradient and from each fraction. The strong enrichment of Fpn and raft-associated proteins in fraction 1 supported its selection for downstream proteomic analysis.

Quantitative proteomic analysis of the fraction 1 clearly indicated that iron overload strongly modifies the protein composition of macrophage membrane rafts. The quality (iTRAQ labeling efficiency reached 99.6%) and coverage (823 proteins detected with greater than 99% confidence) of the proteomic data support the robustness of this analysis and allow a global view of “iron-regulated” (directly or indirectly) proteins in these membrane microdomains. Almost 50% of the total spectra were identified. Iron treatment induced major changes in protein abundance, with high number of proteins being upregulated or downregulated. A high proportion of detected proteins are known to associate with the plasma membrane or lipid rafts (i.e. flotilin1 and 2), suggesting a good quality and purity of the DRM sample. Together, this protein profile of iron-treated macrophage rafts constitute a useful tool for dissecting the molecular mechanisms underlying iron handling, homeostasis, and signaling in macrophages after iron treatment. Notably, our analyses revealed the upregulation of kinases (Taok3, Skap2: ratio =21), phosphatases (Ptpn6: ratio = 16), and other signaling proteins Fyb (ratio =16). The amount of RNF149, an E3 ubiquitin ligase, was also markedly increased (ratio = 9) following iron treatment. Interestingly, another E3 ubiquitin ligase RNF217 is involved in the degradation of Fpn (Jiang et al., 2021). The presence of RNF149 in the microdomain containing Fpn need further investigation.

In addition, some contamination during sub-fractionation cannot be excluded, as no organelle can be purified to complete homogeneity (Zheng & Foster, 2009). The unexpected detection of Fth (iron storage) in our DRM fractions likely reflects co-purification of highly upregulated cytosolic ferritin rather than true lipid raft localization, since FtH is primarily cytosolic and not reported at the cell surface (this study). Similarly, the presence of mitochondrial proteins is consistent with previous DRM proteomic studies and may arise from MAM tethering and/or density overlap with raft fractions (Poston et al., 2011).

### A raft platform against iron mediated oxidative stress

One major observation in our microdomains proteomic analysis is the strong induction of proteins involved in antioxidant defense, such as Gclm, Prdx1, and G6pdx. Such proteins represent complementary components of a cellular antioxidant network, supporting glutathione synthesis, peroxide detoxification, and NADPH production, respectively (Lu, 2013; Perkins et al., 2015). Despite Prxd1 was shown to be present in DRM in different cell types (Woo et al., 2010), direct evidence for iron-induced recruitment of antioxidant enzymes such as Prdx1 or G6pdx to rafts remains limited. Consistent with this possibility, preliminary experiments in FeNTA-treated BMDM showed a redistribution of Prdx1 toward DRM fractions, where it co-fractionated with Hmox1, caveolin1, and flotillin1 (data not shown). Together, Gclm, Prdx1, and G6pdx help to maintain redox homeostasis and limit oxidative damage (i.e. fenton reaction) and lipid peroxidation. Interestingly, lipid rafts and caveolae, a subset of lipid microdomain have been described as key modulators of redox signaling and anti-oxydant response (Li & Gulbins, 2007). Caveolae are small membrane invaginations at the cell surface are formed by caveolins and cavins proteins (Parton et al., 2020). These specific pits at the plasma membrane are described to function in numerous biological processes in response to stimuli (Lamaze et al., 2026). Oxidative stress has been shown to promote the enlargement of membrane microdomains through lipid peroxidation (Ayuyan & Cohen, 2006). Phospholipid oxidation can alter membrane organization and promote either the formation or disruption of lipid rafts. These changes may affect the recruitment of raft-associated proteins, thereby modifying cell signaling and homeostasis. Therefore, the physiological consequences of the membrane reorganizations after iron overload would be a local protection against the iron mediated oxidative stress and in particular against the lipid peroxidation at the plasma membrane.

In addition to these well-known antioxidant proteins, we observed a strong upregulation of Fpn and Hmox1 in DRM of macrophages. Hmox1 is a canonical antioxidant protein through its catabolic activity of heme, which is converted into biliverdin (further reduce to bilirubin, a potent ROS scavenger), CO (anti-oxidant and anti-inflammatory signaling) and Fe²⁺. As an iron exporter, Fpn could be also considered in some extent as an antioxidant protein, protecting the cells from an excess of iron to limit Fenton reaction and ROS production. Interestingly, in addition to its antioxidant function in H₂O₂ detoxification, Prdx1 shares 97% amino acid identity with Hbp23, a heme-binding protein. Prdx1 could therefore play a dual protective role in response to iron/heme-induced stress, either by scavenging excess intracellular heme or through its peroxidase activity by reducing H₂O₂. The detection of both Prdx1/Hbp23 and Hmox1 within the same lipid raft fractions may have biological significance and warrants further investigation.

Our bioinformatic analysis with Ingenuity identified Nrf2 as a potential central regulator of the iron-induced oxidative stress response, a prediction supported by experimental data. Indeed, iron treatment induced nuclear translocation of Nrf2, confirming the activation of the pathway. Prxd1, which is strongly upregulated after iron treatment, is described as an Nrf2-regulated antioxidant genes as well as GCLM and G6pdxx (Kim et al., 2007). In macrophage, Nrf2 is a master regulator of anti-oxidative responses (Ishii et al., 2000; Vomund et al., 2017; Wang et al., 2019). Our functional studies revealed a strict requirement for Nrf2 in basal and iron-induced Fpn expression, as both protein and mRNA induction in these conditions were abolished in Nrf2-deficient macrophages. A strong transcriptional upregulation of Fpn dependent on Nrf2 has been previously described in macrophages treated with electrophilic compounds, heme or oxLDL (Harada et al., 2011; Marques et al., 2016; Marro et al., 2010). The basal expression of Fpn seems to be strongly dependent on Nrf2 [this study and (Harada et al., 2011; Marques et al., 2016)]. Hmox1 is also described as an Nrf2 gene target (Ishii et al., 2000). However, the basal mRNA and protein expression of Hmox1 were not or partially affected in Nrf2 KO BMDM (Marques et al., 2016). In human hepatoma cells, the silencing of Nrf2 with siRNA did not modify basal expression of Hmox1 (Hou et al., 2009). In this study, FeNTA was shown to increase *Hmox1* mRNA in a Nrf2 dependent manner in hepatoma cells. In addition to Nrf2, NFκB and JNK kinases signaling pathways may contribute to *Hmox1* transcription in response to iron-induced oxidative and inflammatory stress (Pronk et al., 2014). Such regulations could explain the maintenance of *Hmox1* mRNA levels in Nrf2-deficient macrophages. Surprisingly, despite a totally blunt mRNA response, Hmox1 protein expression was increased in Nrf2 KO macrophages in response to iron, suggesting the involvement of post-transcriptional regulatory mechanisms. *Hmox1* mRNA contains AU-rich elements (AREs) within its 3′ untranslated region, which are recognized by RNA-binding proteins such as tristetraprolin (TTP) or AUF1 (Choi & Alam, 1996). Binding of these factors can promote mRNA destabilization under basal conditions. Iron or heme overload, through ROS production, may activate p38 MAPK signaling, leading to phosphorylation and displacement of these RNA-binding proteins, thereby allowing translation and rapid accumulation of Hmox1 protein independently of new transcription.

Altogether, our results support the idea that lipid rafts function as membrane platforms organized to prevent oxidative stress–related pathways in macrophages.

### Ferroportin and Hmox1, two partners in the same microdomain

Since the first description of Hmox1 in microsomal fraction of the endoplasmic reticulum (Yoshinaga et al., 1982), the enzyme has been found in many other cellular compartments including mitochondria and nucleus (Dunn et al., 2014). Indeed, Hmox1 belong to the family of tail-anchored proteins, a specific group of proteins present in the cytosol and that can be target and integrate into different cellular membranes via its carboxy-terminal transmembrane domain (Borgese et al., 2003). In our experiment, Hmox1 was detected as two immunoreactive species by western blot in J774 macrophages and to a lesser extent in BMDM. The upper band likely corresponds to full-length Hmox1 (32 kDa) while the lower band may reflect a processed or truncated form of Hmox1. In BMDM, this shorter protein specie was strongly upregulated in NDRM after heme treatment. Interestingly, a faster migrating Hmox1 specie was shown to be enriched in nuclear extracts after heme treatment (Lin et al., 2007). However, additional experiments are required to determine whether it corresponds to a nuclear species in our experiments.

On the other hand, our study demonstrates the presence of full-length Hmox1 in lipid rafts of macrophages after heme (normosang) or iron treatment. A similar observation was shown in rat pulmonary artery endothelial cells after hemin treatment or hypoxia (Kim et al., 2004). The trafficking of Hmox1 into caveolae microdomains has been also reported after LPS or Hypoxia (Kim et al., 2004; Wang et al., 2009). As an example, in LPS treated peritoneal macrophages, the recruitment of Hmox1 and its colocalization/interaction with caveolin1 interfere with the TLR4 proinflammatory signaling (Wang et al., 2009). In our study, both Hmox1 and Fpn were found in caveolae (colocalization with caveolin1) at the cell surface of macrophages, a localization that could be driven by a direct interaction with caveolin1. Interestingly, in J774a1, caveolin1 is not expressed, while caveolin-2 is present at low level and mainly localizes to the Golgi rather than forming clear caveolae at the plasma membrane (Gargalovic & Dory, 2001). However, these macrophages express flotillin-1 and flotillin-2 and may contain flotillin-enriched membrane domains resembling caveolin microdomains. Indeed, flotillin is considered a scaffolding component of lipid rafts that contributes to the partitioning of specific proteins into membrane microdomains during cellular activation (Zhao et al., 2011). It is therefore possible to speculate that in J774a1, in absence of caveolin, Hmox1 is recruited in a flotillin associated raft.

In contrast to Fpn, which is detected in DRM under both basal (despite low) and iron-treated conditions, Hmox1 was mainly associated with non-DRM fractions under basal conditions. Its presence in lipid rafts DRM was markedly induced after iron or heme treatment. Similarly to LPS stimulation (Kim et al., 2004; Wang et al., 2009), the recruitment of Hmox1 to microdomains at the cell surface may result mainly from the targeting of newly synthesized Hmox1 following iron or heme stimulation and need to be explored. In particular, the role of the carboxy-terminal domain in such recruitment process needs to be studied.

### A lipid raft compartment for heme catabolism and iron export

In front of a major cellular uptake of iron (strong erythrophagocytosis activity or heme uptake), the targeting of Hmox1 in the plasma membrane to join Fpn in order to quickly eliminate heme (catabolism) and iron (export) could be a protecting mechanism to prevent iron accumulation in the cells and limit lipid peroxidation in the membrane as discussed before.

The putative degradation of heme and the subsequent iron export in a specific lipid raft structure such as the caveolae is also supported by other observations. Indeed, we previously identified the glycosylphosphatidylinositol (GPI)-anchored ceruloplasmin as a putative functional partner of Fpn within lipid rafts at the macrophage surface (Marques et al., 2012). The GPI-anchored ceruloplasmin was shown to partially co-localized with Fpn in cell surface microdomains of iron-treated BMDM. GPI proteins are described to be enriched in lipid raft and in particular in caveolae (Mayor et al., 1994). At the cell surface, GPI-Cp oxidize the transported ferrous iron (Fe^2+^) to ferric iron (Fe^3+^), a non-toxic form of iron, which is then deliver to the transferrin for subsequent delivery to cells in the body. Beside the GPI-Cp oxidase, the biliverdin reductase (BVR) was also shown to be present in a lipid raft compartment corresponding to caveolae (H. P. Kim et al., 2004; X. M. Wang et al., 2009). The activity of Hmox1 is tightly linked to BVR forming a sequential enzymatic pathway for heme catabolism. The degradation of heme by Hmox1 produce iron, CO and biliverdin that is immediate reduce into bilirubin by BVR. The presence of Cp, BVR, Fpn and Hmox1 in the same compartment at the cell surface is consistent with a coordinated heme degradation and iron efflux at the cell surface in case of an intracellular excess of heme or iron. Iron acts as a key driver of ferroptosis, an iron-dependent form of regulated cell death, by promoting reactive oxygen species generation and iron-dependent phospholipid peroxidation (X. Jiang et al., 2021). Therefore, a lipid raft platform organizes to catabolize heme and export iron from the cell could be considered as a quick and safe way to deplete excess of iron in the cytosol in order to limit the process of ferroptosis and protect the cell from death.

### Conclusions and perspectives

Together, our proteomic results provide a strong rationale for identifying additional Fpn-interacting proteins such as Hmox1 within membrane lipid rafts, especially in caveolae. It suggests that such compartment at the cell surface contribute to the spatial organization and regulation of Fpn in cellular response. It is tempting to speculate that, the heme catabolism with the subsequent iron export by Fpn at the cell surface is a safety way to eliminate iron and limit phospholipids peroxidation (Graphical abstract). Our quantitative proteomic analysis of macrophage lipid rafts also helped to define the local molecular response at the cell surface to cellular stress induced by iron or heme. In such condition, plasma membrane is composed with lipid raft platforms enriched in antioxidant proteins in order to limit the lipid oxidation at that site. Nevertheless, it would be relevant to investigate whether similar changes occur during the process of erythrophagocytosis and heme iron recycling in our macrophages cultures as well as in human and mouse primary macrophages, in particular in splenic red pulp macrophages. Relevant experimental models could also include model with high turnover and recycling of RBC such as transfusion-based erythrophagocytosis models (Youssef et al., 2018), polycythemic mice (Bogdanova et al., 2007), or models of chronic hemolytic anemia, such as pyruvate kinase-deficient mice (Roy et al., 2007).

Overall, the iron-dependent remodeling of the macrophage raft proteome observed here offers a dataset that can be used to uncover new regulators of oxidative stress, iron efflux and homeostasis in macrophages.

## AUTHOR CONTRIBUTIONS

A.W., A.A., and L.R. performed cell culture and iodixanol gradient experiments and western blot. A.W. and A.A. conducted ICW and imaging analyses. A.A. performed immunofluorescence and biotinylation assays. L.R. managed NRF2 knockout mouse breeding. C.F. provided technical assistance for optimization of protein sample preparation and iodixanol gradient analysis, and generate ProteinPilot 3 data processing and Ingenuity Pathway Analysis of MS/MS-identified proteins. M.L. contributed to the generation of proteomic data, including mass spectrometry analyses. L.C. performed proteomic sample preparation, mass spectrometry analysis and participated in manuscript editing. FCH, design experiments, analyzed data, and wrote the manuscript. All authors contributed to data analysis, interpretation, and figure preparation.

## FUNDINGS

This work was supported by INSERM (Institut National de la Santé et de la Recherche Médicale), CNRS (Centre National de la Recherche Scientifique) and ANR (Agence Nationale de la Recherche; ANR-08-GENO-014-03).

## Notes

### Competing Interest Statement

The authors have declared no competing interest.

## References

Abboud, S., & Haile, D. J. (2000). A novel mammalian iron-regulated protein involved in intracellular iron metabolism. The Journal of Biological Chemistry, 275(26), 19906–19912. 10.1074/jbc.M000713200

Auriac, A., Willemetz, A., & Canonne-Hergaux, F. (2010). Lipid raft-dependent endocytosis : A new route for hepcidin-mediated regulation of ferroportin in macrophages. Haematologica, 95(8), 1269–1277. 10.3324/haematol.2009.019992

Ayuyan, A. G., & Cohen, F. S. (2006). Lipid Peroxides Promote Large Rafts : Effects of Excitation of Probes in Fluorescence Microscopy and Electrochemical Reactions during Vesicle Formation. Biophysical Journal, 91(6), 2172–2183. 10.1529/biophysj.106.087387

Besson, C., Willemetz, A., Latour, C., Robert, L., Coppin, H., Roth, M.-P., & Canonne-Hergaux, F. (2024). NEW INSIGHTS INTO THE HEPATIC IRON PHENOTYPE OF BMP6 KNOCKOUT MICE. 10.1101/2023.09.28.559941

Bogdanova, A., Mihov, D., Lutz, H., Saam, B., Gassmann, M., & Vogel, J. (2007). Enhanced erythrophagocytosis in polycythemic mice overexpressing erythropoietin. Blood, 110(2), 762–769. 10.1182/blood-2006-12-063602

Borgese, N., Colombo, S., & Pedrazzini, E. (2003). The tale of tail-anchored proteins : Coming from the cytosol and looking for a membrane. The Journal of Cell Biology, 161(6), 1013–1019. 10.1083/jcb.200303069

Bossart, J., Rippl, A., Barton Alston, A. E., Flühmann, B., Digigow, R., Buljan, M., Ayala-Nunez, V., & Wick, P. (2023). Uncovering the dynamics of cellular responses induced by iron-carbohydrate complexes in human macrophages using quantitative proteomics and phosphoproteomics. Biomedicine & Pharmacotherapy = Biomedecine & Pharmacotherapie, 166, 115404. 10.1016/j.biopha.2023.115404

Canonne-Hergaux, F., Donovan, A., Delaby, C., Wang, H., & Gros, P. (2006). Comparative studies of duodenal and macrophage ferroportin proteins. American Journal of Physiology. Gastrointestinal and Liver Physiology, 290(1), G156–163. 10.1152/ajpgi.00227.2005

Chaston, T., Chung, B., Mascarenhas, M., Marks, J., Patel, B., Srai, S. K., & Sharp, P. (2008). Evidence for differential effects of hepcidin in macrophages and intestinal epithelial cells. Gut, 57(3), 374–382. 10.1136/gut.2007.131722

Choi, A. M., & Alam, J. (1996). Heme Oxygenase-1 : Function, Regulation, and Implication of a Novel Stress-Inducible Protein in Oxidant-Induced Lung Injury. American Journal of Respiratory Cell and Molecular Biology, 15(1), 9–19. 10.1165/ajrcmb.15.1.8679227

Debbiche, R., Ka, C., Gourlaouen, I., Maestri, S., Uguen, K., Jaffrès, P.-A., Callebaut, I., & Le Gac, G. (2023). Cholesterol modulates the human FPN1 iron export function in plasma membrane liquid-ordered microdomains. Biochemistry. 10.1101/2023.12.14.571614

Delaby, C., Pilard, N., Gonçalves, A. S., Beaumont, C., & Canonne-Hergaux, F. (2005). Presence of the iron exporter ferroportin at the plasma membrane of macrophages is enhanced by iron loading and down-regulated by hepcidin. Blood, 106(12), 3979–3984. 10.1182/blood-2005-06-2398

Delaby, C., Pilard, N., Hetet, G., Driss, F., Grandchamp, B., Beaumont, C., & Canonne-Hergaux, F. (2005). A physiological model to study iron recycling in macrophages. Experimental Cell Research, 310(1), 43–53. 10.1016/j.yexcr.2005.07.002

Dhungana, S., Merrick, B. A., Tomer, K. B., & Fessler, M. B. (2009). Quantitative proteomics analysis of macrophage rafts reveals compartmentalized activation of the proteasome and of proteasome-mediated ERK activation in response to lipopolysaccharide. Molecular & Cellular Proteomics: MCP, 8(1), 201–213. 10.1074/mcp.M800286-MCP200

Donovan, A., Brownlie, A., Zhou, Y., Shepard, J., Pratt, S. J., Moynihan, J., Paw, B. H., Drejer, A., Barut, B., Zapata, A., Law, T. C., Brugnara, C., Lux, S. E., Pinkus, G. S., Pinkus, J. L., Kingsley, P. D., Palis, J., Fleming, M. D., Andrews, N. C., & Zon, L. I. (2000). Positional cloning of zebrafish ferroportin1 identifies a conserved vertebrate iron exporter. Nature, 403(6771), 776–781. 10.1038/35001596

Dunn, L. L., Midwinter, R. G., Ni, J., Hamid, H. A., Parish, C. R., & Stocker, R. (2014). New insights into intracellular locations and functions of heme oxygenase-1. Antioxidants & Redox Signaling, 20(11), 1723–1742. 10.1089/ars.2013.5675

Frazer, D. M., Wilkins, S. J., Darshan, D., Mirciov, C. S. G., Dunn, L. A., & Anderson, G. J. (2017). Ferroportin Is Essential for Iron Absorption During Suckling, But Is Hyporesponsive to the Regulatory Hormone Hepcidin. Cellular and Molecular Gastroenterology and Hepatology, 3(3), 410–421. 10.1016/j.jcmgh.2016.12.002

Gargalovic, P., & Dory, L. (2001). Caveolin1 and caveolin-2 expression in mouse macrophages. High density lipoprotein 3-stimulated secretion and a lack of significant subcellular co-localization. The Journal of Biological Chemistry, 276(28), 26164–26170. 10.1074/jbc.M011291200

Harada, N., Kanayama, M., Maruyama, A., Yoshida, A., Tazumi, K., Hosoya, T., Mimura, J., Toki, T., Maher, J. M., Yamamoto, M., & Itoh, K. (2011). Nrf2 regulates ferroportin 1-mediated iron efflux and counteracts lipopolysaccharide-induced ferroportin 1 mRNA suppression in macrophages. Archives of Biochemistry and Biophysics, 508(1), 101–109. 10.1016/j.abb.2011.02.001

Hou, W.-H., Rossi, L., Shan, Y., Zheng, J.-Y., Lambrecht, R.-W., & Bonkovsky, H.-L. (2009). Iron increases HMOX1 and decreases hepatitis C viral expression in HCV-expressing cells. World Journal of Gastroenterology, 15(36), 4499–4510. 10.3748/wjg.15.4499

Ishii, T., Itoh, K., Takahashi, S., Sato, H., Yanagawa, T., Katoh, Y., Bannai, S., & Yamamoto, M. (2000). Transcription factor Nrf2 coordinately regulates a group of oxidative stress-inducible genes in macrophages. The Journal of Biological Chemistry, 275(21), 16023–16029. 10.1074/jbc.275.21.16023

Jiang, L., Wang, J., Wang, K., Wang, H., Wu, Q., Yang, C., Yu, Y., Ni, P., Zhong, Y., Song, Z., Xie, E., Hu, R., Min, J., & Wang, F. (2021). RNF217 regulates iron homeostasis through its E3 ubiquitin ligase activity by modulating ferroportin degradation. Blood, 138(8), 689–705. 10.1182/blood.2020008986

Jiang, X., Stockwell, B. R., & Conrad, M. (2021). Ferroptosis : Mechanisms, biology and role in disease. Nature Reviews. Molecular Cell Biology, 22(4), 266–282. 10.1038/s41580-020-00324-8

Kim, H. P., Wang, X., Galbiati, F., Ryter, S. W., & Choi, A. M. K. (2004). Caveolae compartmentalization of heme oxygenase-1 in endothelial cells. FASEB Journal: Official Publication of the Federation of American Societies for Experimental Biology, 18(10), 1080–1089. 10.1096/fj.03-1391com

Kim, Y.-J., Ahn, J.-Y., Liang, P., Ip, C., Zhang, Y., & Park, Y.-M. (2007). Human prx1 gene is a target of Nrf2 and is up-regulated by hypoxia/reoxygenation : Implication to tumor biology. Cancer Research, 67(2), 546–554. 10.1158/0008-5472.CAN-06-2401

Knutson, M. D., Oukka, M., Koss, L. M., Aydemir, F., & Wessling-Resnick, M. (2005). Iron release from macrophages after erythrophagocytosis is up-regulated by ferroportin 1 overexpression and down-regulated by hepcidin. Proceedings of the National Academy of Sciences of the United States of America, 102(5), 1324–1328. 10.1073/pnas.0409409102

Knutson, M. D., Vafa, M. R., Haile, D. J., & Wessling-Resnick, M. (2003). Iron loading and erythrophagocytosis increase ferroportin 1 (FPN1) expression in J774 macrophages. Blood, 102(12), 4191–4197. 10.1182/blood-2003-04-1250

Lamaze, C., Blouin, C. M., & Sens, P. (2026). Caveolae mechanics in cellular functions and disease. Nature Reviews Molecular Cell Biology, 27(8), 581–600. 10.1038/s41580-026-00964-2

Li, N., Mak, A., Richards, D. P., Naber, C., Keller, B. O., Li, L., & Shaw, A. R. E. (2003). Monocyte lipid rafts contain proteins implicated in vesicular trafficking and phagosome formation. Proteomics, 3(4), 536–548. 10.1002/pmic.200390067

Li, P.-L., & Gulbins, E. (2007). Lipid Rafts and Redox Signaling. Antioxidants & Redox Signaling, 9(9), 1411–1416. 10.1089/ars.2007.1736

Lin, Q., Weis, S., Yang, G., Weng, Y.-H., Helston, R., Rish, K., Smith, A., Bordner, J., Polte, T., Gaunitz, F., & Dennery, P. A. (2007). Heme Oxygenase-1 Protein Localizes to the Nucleus and Activates Transcription Factors Important in Oxidative Stress. Journal of Biological Chemistry, 282(28), 20621–20633. 10.1074/jbc.M607954200

Lingwood, D., & Simons, K. (2010). Lipid rafts as a membrane-organizing principle. *Science (New York*, N.Y.), 327(5961), 46–50. 10.1126/science.1174621

Lu, S. C. (2013). Glutathione synthesis. Biochimica et Biophysica Acta (BBA) - General Subjects, 1830(5), 3143–3153. 10.1016/j.bbagen.2012.09.008

Marques, L., Auriac, A., Willemetz, A., Banha, J., Silva, B., Canonne-Hergaux, F., & Costa, L. (2012). Immune cells and hepatocytes express glycosylphosphatidylinositol-anchored ceruloplasmin at their cell surface. Blood Cells, Molecules & Diseases, 48(2), 110–120. 10.1016/j.bcmd.2011.11.005

Marques, L., Negre-Salvayre, A., Costa, L., & Canonne-Hergaux, F. (2016). Iron gene expression profile in atherogenic Mox macrophages. Biochimica Et Biophysica Acta, 1862(6), 1137–1146. 10.1016/j.bbadis.2016.03.004

Marro, S., Chiabrando, D., Messana, E., Stolte, J., Turco, E., Tolosano, E., & Muckenthaler, M. U. (2010). Heme controls ferroportin1 (FPN1) transcription involving Bach1, Nrf2 and a MARE/ARE sequence motif at position-7007 of the FPN1 promoter. Haematologica, 95(8), 1261–1268. 10.3324/haematol.2009.020123

Mayor, S., Rothberg, K. G., & Maxfield, F. R. (1994). Sequestration of GPI-anchored proteins in caveolae triggered by cross-linking. Science, 264(5167), 1948–1951. 10.1126/science.7516582

McKie, A. T., Marciani, P., Rolfs, A., Brennan, K., Wehr, K., Barrow, D., Miret, S., Bomford, A., Peters, T. J., Farzaneh, F., Hediger, M. A., Hentze, M. W., & Simpson, R. J. (2000). A novel duodenal iron-regulated transporter, IREG1, implicated in the basolateral transfer of iron to the circulation. Molecular Cell, 5(2), 299–309. 10.1016/s1097-2765(00)80425-6

Muckenthaler, M. U., Galy, B., & Hentze, M. W. (2008). Systemic iron homeostasis and the iron-responsive element/iron-regulatory protein (IRE/IRP) regulatory network. Annual Review of Nutrition, 28, 197–213. 10.1146/annurev.nutr.28.061807.155521

Nemeth, E., Tuttle, M. S., Powelson, J., Vaughn, M. B., Donovan, A., Ward, D. M., Ganz, T., & Kaplan, J. (2004). Hepcidin regulates cellular iron efflux by binding to ferroportin and inducing its internalization. Science, 306(5704), 2090–2093. 10.1126/science.1104742

Parton, R. G., Kozlov, M. M., & Ariotti, N. (2020). Caveolae and lipid sorting : Shaping the cellular response to stress. Journal of Cell Biology, 219(4), e201905071. 10.1083/jcb.201905071

Perkins, A., Nelson, K. J., Parsonage, D., Poole, L. B., & Karplus, P. A. (2015). Peroxiredoxins : Guardians against oxidative stress and modulators of peroxide signaling. Trends in Biochemical Sciences, 40(8), 435–445. 10.1016/j.tibs.2015.05.001

Polati, R., Castagna, A., Bossi, A. M., Alberio, T., De Domenico, I., Kaplan, J., Timperio, A. M., Zolla, L., Gevi, F., D’Alessandro, A., Brunch, R., Olivieri, O., & Girelli, D. (2012). Murine macrophages response to iron. Journal of Proteomics, 76 Spec No., 10–27. 10.1016/j.jprot.2012.07.018

Poston, C. N., Duong, E., Cao, Y., & Bazemore-Walker, C. R. (2011). Proteomic analysis of lipid raft-enriched membranes isolated from internal organelles. Biochemical and Biophysical Research Communications, 415(2), 355–360. 10.1016/j.bbrc.2011.10.072

Pronk, T. E., van der Veen, J. W., Vandebriel, R. J., van Loveren, H., de Vink, E. P., & Pennings, J. L. A. (2014). Comparison of the molecular topologies of stress-activated transcription factors HSF1, AP-1, NRF2, and NF-κB in their induction kinetics of HMOX1. Bio Systems, 124, 75–85. 10.1016/j.biosystems.2014.09.005

Roy, M.-F., Riendeau, N., Bédard, C., Hélie, P., Min-Oo, G., Turcotte, K., Gros, P., Canonne-Hergaux, F., & Malo, D. (2007). Pyruvate kinase deficiency confers susceptibility to Salmonella typhimurium infection in mice. The Journal of Experimental Medicine, 204(12), 2949–2961. 10.1084/jem.20062606

Sabelli, M., Montosi, G., Garuti, C., Caleffi, A., Oliveto, S., Biffo, S., & Pietrangelo, A. (2017). Human macrophage ferroportin biology and the basis for the ferroportin disease. Hepatology, 65(5), 1512–1525. 10.1002/hep.29007

Schuck, S., Honsho, M., Ekroos, K., Shevchenko, A., & Simons, K. (2003). Resistance of cell membranes to different detergents. Proceedings of the National Academy of Sciences, 100(10), 5795–5800. 10.1073/pnas.0631579100

Simons, K., & Sampaio, J. L. (2011). Membrane organization and lipid rafts. Cold Spring Harbor Perspectives in Biology, 3(10), a004697. 10.1101/cshperspect.a004697

Soares, M. P., & Hamza, I. (2016). Macrophages and Iron Metabolism. Immunity, 44(3), 492–504. 10.1016/j.immuni.2016.02.016

Vomund, S., Schäfer, A., Parnham, M., Brüne, B., & Von Knethen, A. (2017). Nrf2, the Master Regulator of Anti-Oxidative Responses. International Journal of Molecular Sciences, 18(12), 2772. 10.3390/ijms18122772

Wang, P., Geng, J., Gao, J., Zhao, H., Li, J., Shi, Y., Yang, B., Xiao, C., Linghu, Y., Sun, X., Chen, X., Hong, L., Qin, F., Li, X., Yu, J.-S., You, H., Yuan, Z., Zhou, D., Johnson, R. L., & Chen, L. (2019). Macrophage achieves self-protection against oxidative stress-induced ageing through the Mst-Nrf2 axis. Nature Communications, 10(1), 755. 10.1038/s41467-019-08680-6

Wang, X. M., Kim, H. P., Nakahira, K., Ryter, S. W., & Choi, A. M. K. (2009). The Heme Oxygenase-1/Carbon Monoxide Pathway Suppresses TLR4 Signaling by Regulating the Interaction of TLR4 with Caveolin1. The Journal of Immunology, 182(6), 3809–3818. 10.4049/jimmunol.0712437

Woo, H. A., Yim, S. H., Shin, D. H., Kang, D., Yu, D.-Y., & Rhee, S. G. (2010). Inactivation of Peroxiredoxin I by Phosphorylation Allows Localized H2O2 Accumulation for Cell Signaling. Cell, 140(4), 517–528. 10.1016/j.cell.2010.01.009

Yoshinaga, T., Sassa, S., & Kappas, A. (1982). Purification and properties of bovine spleen heme oxygenase. Amino acid composition and sites of action of inhibitors of heme oxidation. The Journal of Biological Chemistry, 257(13), 7778–7785.

Youssef, L. A., Rebbaa, A., Pampou, S., Weisberg, S. P., Stockwell, B. R., Hod, E. A., & Spitalnik, S. L. (2018). Increased erythrophagocytosis induces ferroptosis in red pulp macrophages in a mouse model of transfusion. Blood, 131(23), 2581–2593. 10.1182/blood-2017-12-822619

Zhao, F., Zhang, J., Liu, Y.-S., Li, L., & He, Y.-L. (2011). Research advances on flotillins. Virology Journal, 8(1), 479. 10.1186/1743-422X-8-479

Zheng, Y. Z., & Foster, L. J. (2009). Contributions of quantitative proteomics to understanding membrane microdomains. Journal of Lipid Research, 50(10), 1976–1985. 10.1194/jlr.R900018-JLR200

