## Supplemental Figures for "QUANTITATIVE PROTEOMIC REVEALS HEME OXYGENASE-1 LOCALIZATION TO CELL-SURFACE LIPID RAFTS AND ITS ASSOCIATION WITH FERROPORTIN IN IRON-LOADED MACROPHAGES"

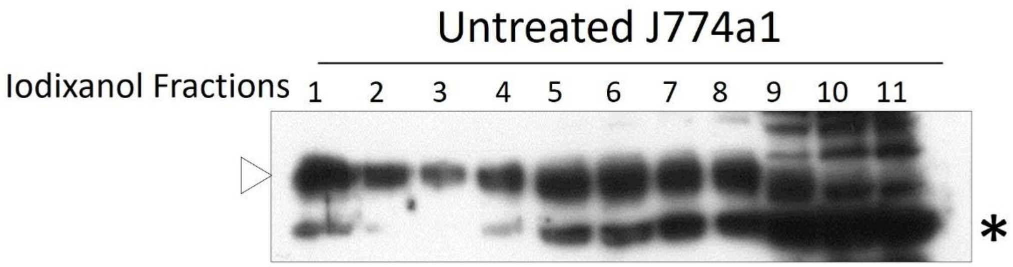


**Fig.S1**. Long exposure of the Fpn signal distribution in iodixanol fractions detected in untreated J774a1 cells. (*) unspecific band.

---

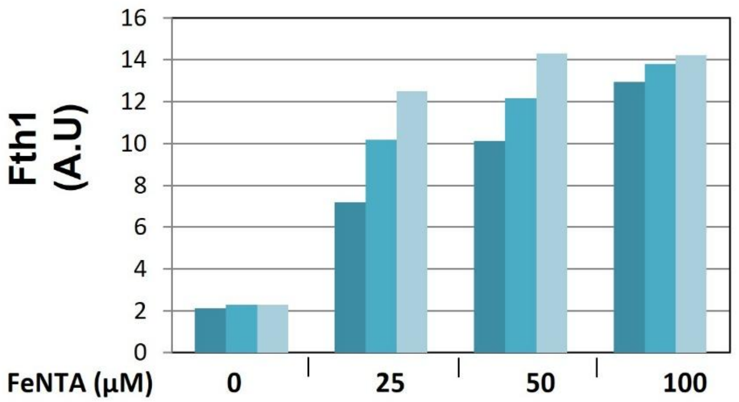


**Fig.S2**: Dose-dependent effects of FeNTA (0–100 µM) on Ferritin H expression measured by In Cell western Blot (3 independent wells)

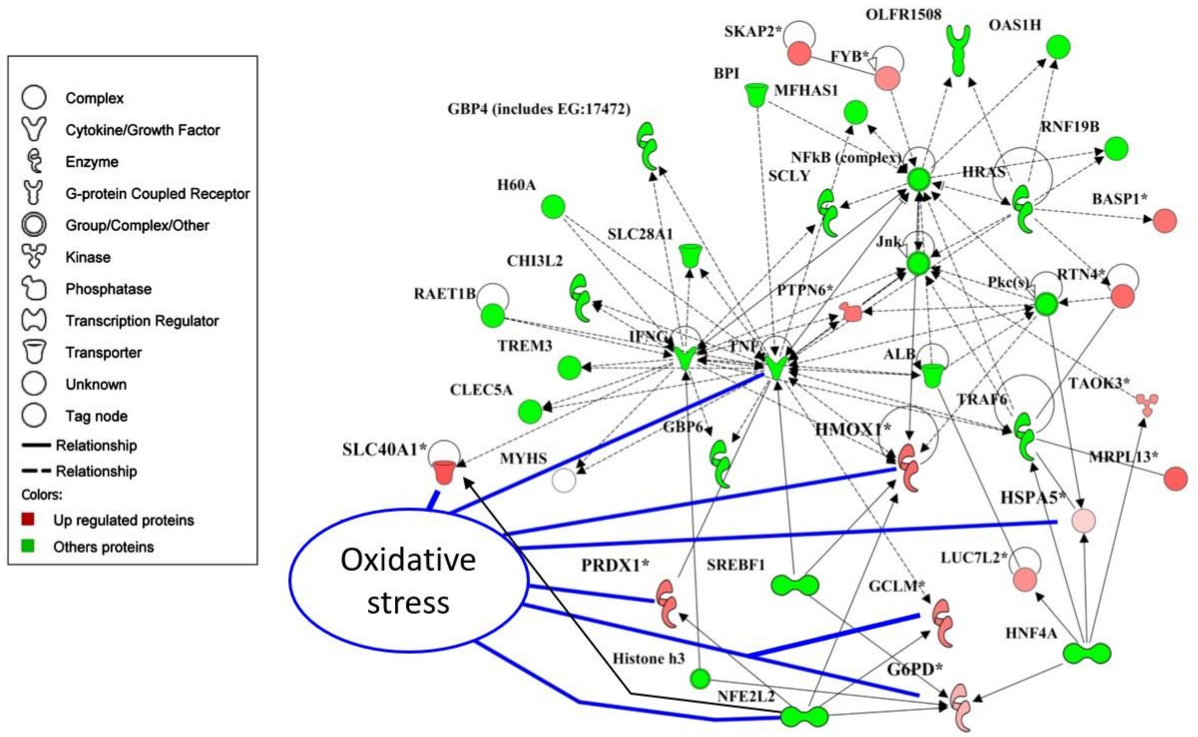


**Fig.S3. Ingenuity Pathway Analysis (IPA) network linking iron-induced proteomic changes to oxidative stress.** Proteins identified by iTRAQ proteomics were mapped using IPA. Node shapes represent functional categories (see legend, left), and edges indicate known or predicted interactions. Red nodes correspond to upregulated proteins in response to iron, while green nodes denote additional putative interacting proteins. The network highlights the upregulation of *HMOX1 (Heme oxygenase1) and SLC40A1 (ferroportin)*, with *PRDX1* (peroxiredoxin 1), *G6PD* (glucose-6-phosphate dehydrogenase), *HSPA5* (heat shock protein A5), and *GCLM (*Glutamate–Cysteine Ligase). Altogether, such regulations support activation of antioxidant and stress-protective pathways, consistent with a cellular adaptation to iron overload. The transcription factor Nrf2 (NFE2L2), positioned at the core of the network, appears as a master regulator driving the coordinated induction of these antioxidant and cytoprotective proteins in response to FeNTA.

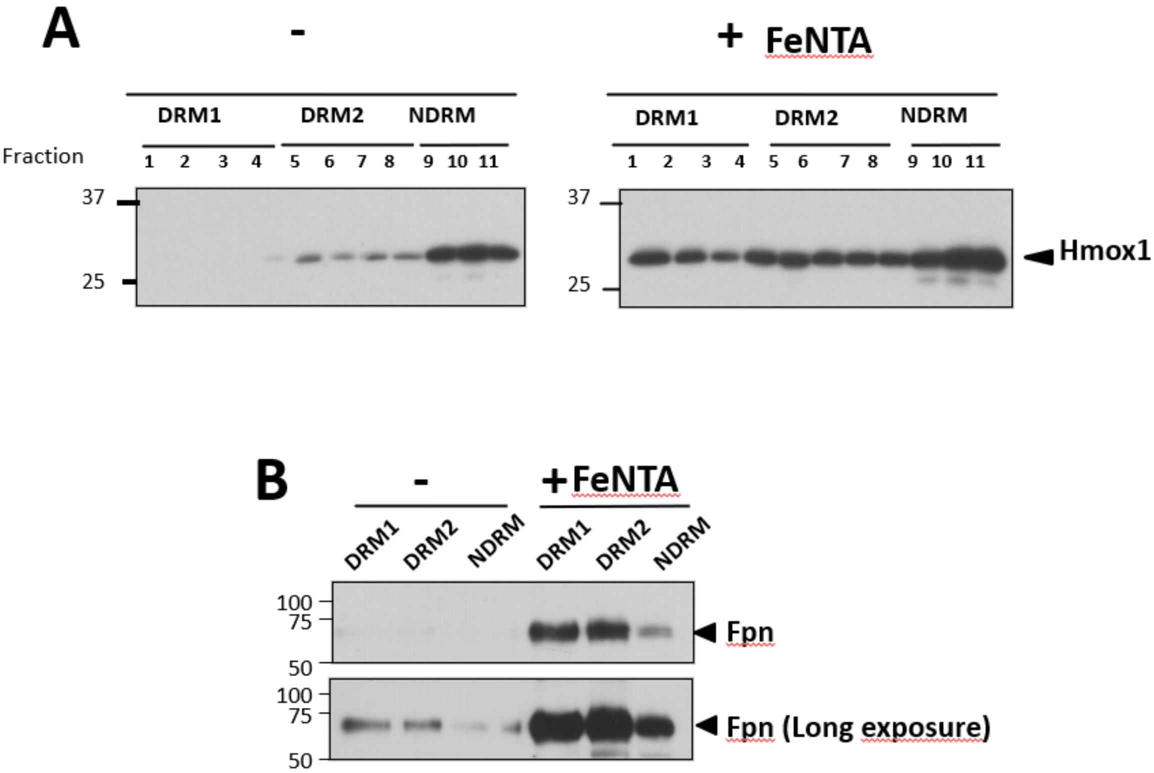


**Fig.S4. (A)** Western blot analysis of heme oxygenase (Hmox1) expression in different protein fractions (1 to 11) obtained from an iodixanol gradient of proteins extracted from BMDM untreated (-) or treated with iron. **(B)** A long exposure of the Fpn detection indicate that in basal conditions the iron exporter is mostly expressed in DRM1 and 2. Molecular weights are shown on the right.

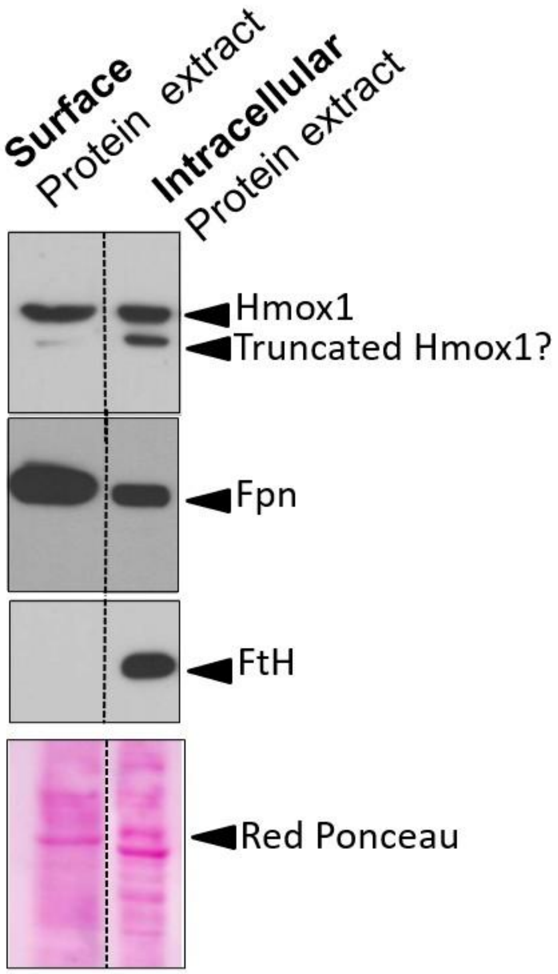


**Fig.S5. Western blot analysis of surface and intracellular proteins following cell surface biotinylation.** After iron treatment**,** cell surface proteins from J774a1 were isolated via streptavidin pull-down, allowing direct comparison of protein expression in surface (biotinylated) versus intracellular (non-biotinylated) fractions. Biotinylation experiment confirmed that Hmox1 is detected with Fpn the cell surface at the plasma membrane.

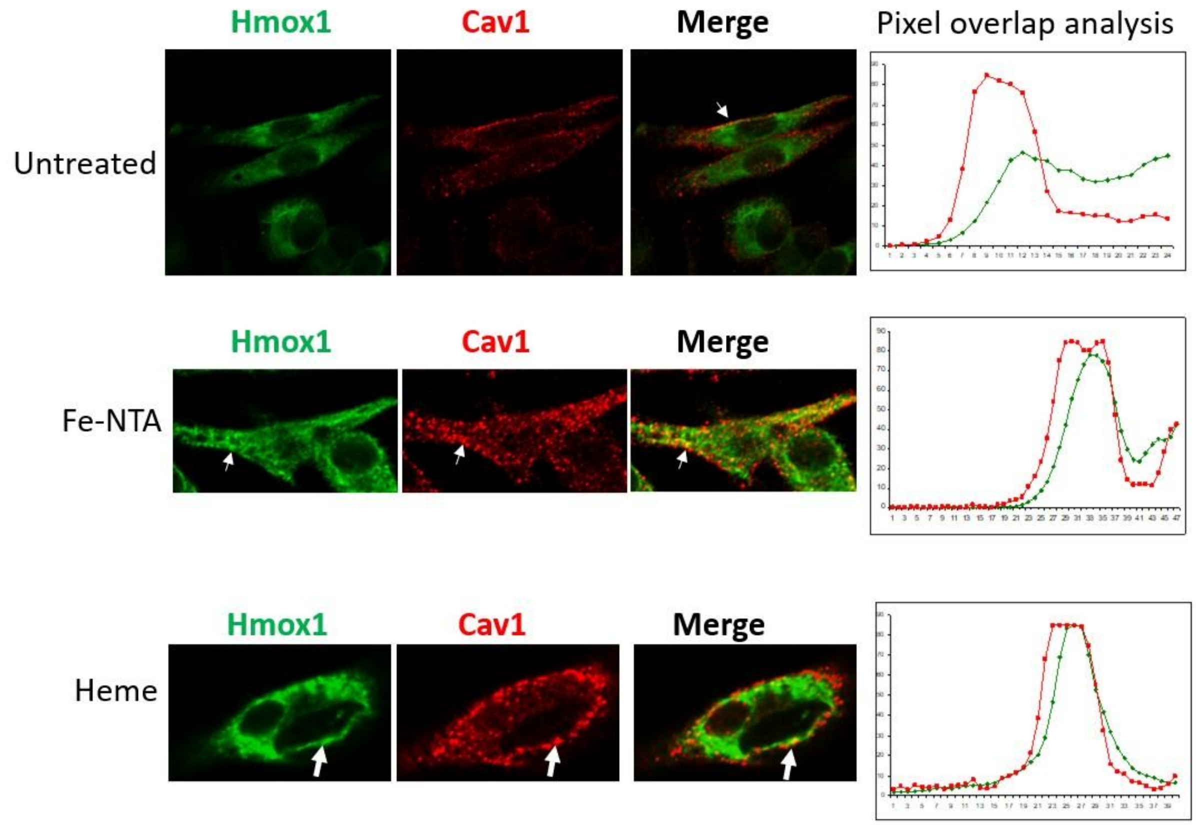


**Fig.S6. Colocalization of Hmox1 with Caveolin-1 (Cav1) in macrophages under iron or heme treatment.** Immunofluorescence analysis of Hmox1 (green) and Caveolin-1 (red) in untreated, FeNTA–treated, or heme-treated macrophages. In untreated cells, Hmox1 shows a diffuse cytoplasmic distribution with no overlap with Caveolin-1–positive domains. Following FeNTA exposure, Hmox1 expression increases and partially colocalizes with Caveolin-1 at discrete membrane-associated sites (arrows). Heme treatment strongly induces Hmox1, with colocalization at the plasma membrane with caveolin1. In untreated cells, pixel intensity profiles (right panels) show minimal overlap between Hmox1 (green) and Caveolin-1 (red) signals, indicating that Hmox1 is largely absent from Caveolin-1–positive membrane domains under basal conditions. On the other hand, the Pixel intensity overlap profiles after FeNTA and heme treatments illustrated increased spatial correlation between Hmox1 (green) and Caveolin-1 (red) signals.
